# Dynamics of the membrane- and cell wall-associated proteome of *Arabidopsis thaliana* roots in response to uranium stress

**DOI:** 10.1101/2024.02.02.578547

**Authors:** Jonathan Przybyla-Toscano, Cherif Chetouhi, Thierry Balliau, Claude Alban, Jacques Bourguignon, Stéphane Ravanel

## Abstract

Uranium (U) is a non-essential and toxic metal for plants, which have the ability to take up uranyl ions from the soil and preferentially accumulate them in the roots. We showed that the bulk of the radionuclide accumulates in the root insoluble proteome of Arabidopsis plants challenged with U. Therefore, to elucidate new molecular mechanisms related to U stress response and tolerance, we used label-free quantitative proteomics to analyze the dynamics of the root membrane- and cell wall-enriched proteome under U stress. Of the 2,802 proteins identified, 458 showed differential accumulation in response to U. Biological processes affected by U include response to stress, amino acid metabolism, and previously unexplored functions associated with membranes and the cell wall. Indeed, our analysis supports a dynamic and complex reorganization of the cell wall in response to U stress, including lignin and suberin synthesis, pectin modifications, polysaccharide hydrolysis, and Casparian strips formation. Water flux through aquaporins and vesicular trafficking were also significantly perturbed by U stress. Finally, the abundance of metal transporters and iron, calcium, and other metal-binding proteins was affected by U. These proteins may play a role in controlling the fate and toxicity of U in plants.

## 1 INTRODUCTION

Heavy metals are by nature present in the earth’s crust composition. In addition, industrial and agricultural activities have direct consequences in their redistribution in the environment. This situation can lead to the accumulation of non-essential trace metals in the soil, and consequently represent a threat to the environment and food safety due to their non-biodegradability, bioavailability and toxicity to crops. This is for instance the case for uranium (U). This actinide element is naturally dispersed in rocks and soils at an average concentration of 1-4 ppm (Anke et al., 2009) and has been found at high concentrations in different area around the world (see Chen et al., 2021 for an overview). As examples, up to 250 mg U.kg^-1^ were detected in sampling sites from Cunha Baixa U mine area in Portugal (Neves and Abreu, 2009) and up to 3500 mg U.kg^-1^ were detected in soils surrounding the reclaimed U mine of Rophin in France (Martin et al., 2020). A field experiment conducted on edible vegetables grown in the agricultural area of Cunha Baixa showed that the amounts of U in the editable tissues of lettuce, potato, green bean, carrot, cabbage, apple and maize were strongly accumulated (Neves and Abreu, 2009; Neves et al., 2012). In agricultural soils, U is also widely dispersed due to the extensive and long-term use of mineral fertilizers and the significant U contamination of phosphate ores (Vandenhove, 2002). In the environment, U coexists as U (+VI) and U (+IV) valence states and the uranyl ions (UO_2_^2+^) are the most abundant form of U in its oxidized state. In this form, uranyl cations react with inorganic anions or organic acids to form highly mobile and soluble complexes in the rhizosphere that are bioavailable and can be absorbed by plants (Hu et al., 2024).

Being sessile, plants have to cope with varying U concentrations in the environment, some of them being detrimental for growth. The chemical toxicity of U in plants has been analyzed in different species. In most species, U is mainly accumulated in roots and its translocation to aerial part is limited (Chen et al., 2021). Analysis of the subcellular distribution of U from roots and protoplasts showed that this radionuclide is primarily sequestrated in the cell wall, whereas only small amounts of cellular U are present in the soluble fraction (Lai et al., 2020; Sarthou et al., 2020; Chen et al., 2023). Uranium accumulation in plant tissues causes an inhibition of plant growth and root elongation, by interfering with carbon and nitrogen assimilation, photosynthesis, mineral nutrition (*i.e.* iron, calcium, phosphorus, magnesium), and hormone synthesis and distribution (*i.e.* auxin, jasmonic acid and salicylic acid) (Vanhoudt et al., 2010; Vanhoudt et al., 2011; Doustaly et al., 2014; Saenen et al., 2014; Saenen et al., 2015; Serre et al., 2019; Wu et al., 2022; Chen et al., 2022; Chen et al., 2023). In addition, U exposure leads to an overproduction of reactive oxygen species (ROS) (Gupta et al., 2020; Wu et al., 2022; Xiao et al., 2023). However, the molecular mechanisms behind these effects are still poorly understood. To date, the pathway for U entry in root cells is the best characterized. It was demonstrated that calcium-permeable channels are required for U uptake (Rajabi et al., 2021; Sarthou et al., 2022), while the main high affinity iron transporter IRT1 is dispensable in Arabidopsis (Berthet et al., 2018). The calcium channel-dependent pathway for U uptake is conserved in yeast (Revel et al., 2022) and could represent a general uptake mechanism, at least in the eukaryotic lineage. In addition, the calcium concentration in the nutrient medium modulates U responses in shoots (Mertens et al., 2022). Beside transporter-mediated uptake, an additional pathway implying endocytic uptake might also be important for U transport into plant cells (John et al., 2022).

During the last decade, several comparative genomic approaches and/or quantitative analyses performed in *A. thaliana*, *Vicia faba* and *Ipomoea batatas* roots have provided insight into global gene expression and metabolic adjustments in response to U (Doustaly et al., 2014; Lai et al., 2020; Sarthou et al., 2020; Zhang et al., 2020; Lai et al., 2021, Vallet et al., 2023; Xiao et al., 2023; Chen et al., 2023). To shed light on the consequences of uranyl ions on the soluble proteome of Arabidopsis root and shoot cells, we previously developed an ionomic and top-down proteomic analysis coupled with biochemical and structural approaches (Sarthou et al., 2020; Vallet et al., 2023). In these studies, we identified 38 proteins able to bind U *in vitro* and we demonstrated that the Arabidopsis cation-binding protein PCaP1 is able to bind U(VI) in addition to other metals (*i.e.* Ca(II), Cu(II) and Fe(III) (Vallet et al., 2023). This newly identified U-binding protein, either associated with the plasma membrane or soluble in the cytosol, was initially described to play a role in calcium signaling, but also to regulate the actin and the microtubule cytoskeleton in a calcium-dependent manner (Nagasaki et al., 2008; Li et al., 2011). While most of U is associated with insoluble cellular fractions, i.e. cell wall, membranes, and high-molecular-weight complexes (Sarthou et al., 2020), our knowledge of the effect of U on these cellular and extracellular structures is very limited. To date, the only documented example of a U-binding membrane protein in any organism is the bacterial UipA protein (Gallois et al., 2022). This single-pass transmembrane protein contains a large domain with nanomolar affinity for uranyl and Fe(III), and is essential for bacterial tolerance to the radionuclide.

In an effort to elucidate the molecular mechanisms of U stress response and tolerance in plants, the aim of this study was to identify membrane- and cell wall-associated proteins in Arabidopsis roots whose expression is regulated upon U stress. To this end, we have developed a label-free quantitative proteomic workflow based on nano-LC-MS/MS analysis followed by a comprehensive computational study. Using this approach, we analyzed the dynamics of the Arabidopsis root membrane- and cell wall-enriched proteome under U stress and identified 458 proteins differentially regulated by U. Our approach targeting the major site of U accumulation in root cells revealed unprecedented biological processes affected by the radionuclide, including cell wall organization, radial apoplastic transport, water flux, and endomembrane trafficking. Our analysis also highlighted potential transporters and metal-binding proteins involved in the fate of U in plants.

## 2 MATERIAL AND METHODS

### 2.1 Plant cultivation and uranium treatment

*Arabidopsis thaliana* Columbia-0 (Col-0) wild type (WT) was grown under hydroponic conditions using the experimental device described in Berthet et al. (2018). Plants were grown under a short-day photoperiod (8 h of light, 80 μmol photons m^-2^ s^-1^ photosynthetically active radiation) at 60% relative humidity and 20°C. Seeds were initially stratified in water for 3 days at 4°C, and then placed in perforated cups filled with 0.5% (w/v) agar. The cups containing seedlings were transferred to floating supports in black polypropylene containers filled with 200 ml of "Gre medium" (0.8 mM K_2_SO_4_, 1 mM Ca(NO_3_)_2_, 1 mM MgSO_4_, 0.25 mM KH_2_PO_4_, 10 µM H_3_BO_3_, 0.2 µM CuSO_4_, 2 µM MnSO_4_, 0.01 µM (NH_4_)_6_Mo_7_O_24_, 2 µM ZnSO_4_, 10 µM NaCl, 0.02 µM CoCl_2_, 20 μM FeNaEDTA), pH 5.6, and cultivated for 4 to 5 weeks. Then, plants were transferred to distilled water supplemented or not with 5 µM or 50 μM uranyl nitrate (UO_2_(NO_3_)_2_) during 48 h. To perform proteomic and ionomic analyses, excess of U was removed from the root surface by a washing step with a carbonate solution (10 mM Na_2_CO_3_,), followed by two additional washes with distilled water. Finally, roots were carefully dried on absorbent paper. Six biological replicates were performed for each concentration (control, 5 and 50 µM of uranyl nitrate). Each replicate is equivalent to the roots of three independent plants.

### 2.2 Uranium quantification by Inductively Coupled Plasma Mass Spectrometry (ICP-MS)

For U determination, digestion of roots was performed at 90°C during 4 h in 400 µL of 65%(w/v) HNO_3_ (Suprapur, Merck). Mineralized samples were diluted in 0.65% (w/v) HNO_3_ and U quantification was performed using an iCAP RQ ICP-MS (Thermo Fisher Scientific GmbH, Germany) operating in standard mode. The concentration was determined using a standard curve and a standard internal solution containing rhodium and ytterbium. The Qtegra software was used for data acquisition and integration.

### 2.3 Protein extraction and quantification

Proteins were extracted from 400 to 700 mg (fresh weight) of roots. Root tissues were ground with liquid nitrogen in a mortar before homogenization with the extraction solution (20 mM Tris-HCl, pH 7.0, 1 mM EDTA, 1 mM DTT and a cocktail of protease inhibitors, Roche). The suspension was ultracentrifuged at 105,000 g for 20 min. The supernatant containing the soluble proteins was collected, while the pellet was washed twice with the extraction solution and then resuspended in the extraction solution supplemented with 1% (w/v) SDS. The suspension was incubated at 4°C for 30 min to allow protein solubilization by SDS and then centrifuged at 15,000 g for 20 min. The supernatant was collected to recover proteins solubilized by SDS. Protein quantification of the extracts was estimated using the BCA Protein Assay Pierce (Thermo Scientific) and bovine serum albumin as standard.

### 2.4 SDS PAGE immunoblot assay

Soluble or SDS-solubilized proteins from root extracts were separated on 12% (w/v) reducing polyacrylamide gels (SDS-PAGE) and transferred onto nitrocellulose membrane. After a blocking step with 4% (w/v) BSA in TBS-0.1% (v/v) Tween, the immunoblot reaction was performed overnight at 4°C with polyclonal antibodies raised against the membrane tonoplast intrinsic protein (TIP) (Agrisera AS09493) or the soluble fructose-bisphosphate aldolases (FBAs) (Mininno et al, 2012). Following three washes, membranes were incubated during 1 h at room temperature with secondary antibodies conjugated to horseradish peroxidase. Signal detection was performed using the ECL prime detection reagent (Amersham) and fluorescence was visualized using ImageQuant 800 (Amersham) imaging system.

### 2.5 Proteomic preparation and label-free nanoLC-MS/MS analysis

About 15 µg of proteins were separated on SDS-PAGE gels (Bio-Rad 3450009) at a constant voltage of 200V for 12 min. Gels were stained with the Bio-Safe™ Coomassie Stain solution (Bio-Rad 1610786) according to the manufacturer recommendations. Proteins were fixed with a 10% (v/v) acetic acid and 40% (v/v) ethanol solution, and rinsed with distilled water. After acquisition of a gel image, the tracks were cut into three bands corresponding to the three protein subfractions. The bands were rinsed and digested with 200 ng trypsin in a final volume of 100 µL. After digestion, samples were dried using a SPD111 SpeedVac (Thermo Scientific) until complete evaporation. LC-MS/MS analyzes were performed using a NanoLC-Ultra system (nano2DUltra, Eksigent) connected to a Q-Exactive plus mass spectrometer (Thermo Electron, Waltham, MA, USA). For each sample, approximately 400 ng of the peptides were loaded onto a Biosphere C18 120 Angstrom precolumn (2 cm, 100 µm, 5 μm; nanoseparation) at 7,500 nL min-1 and desalted with 0.1% (v/v) formic acid and 2% (v/v) acetonitrile. After 5 min, the precolumn was connected to a Biosphere C18 120 Angstrom nanocolumn (30 cm, 75 µm, 3 μm; nanoseparation). The gradient profile contained 5 steps corresponding to: step 0 (0 min in 95% buffer A [0.1% (v/v) formic acid] and 5% buffer B [0.1% (v/v) formic acid in acetonitrile]), step 1 (75 min in 70% buffer A and 30% buffer B), step 2 (80 min in 5% buffer A and 95% buffer B), step 3 (85 min in 5% buffer A and 95% buffer B), step 4 (88 min in 95% buffer A and 5% buffer B) and step 5 (95 min in 95% buffer A and 5% buffer B). Nano-ESI was performed with a spray voltage of 1.6 kV and a heated capillary temperature of 250°C. The acquisition was performed using XCalibur 4.0.29 (Thermo Electron) software in data-dependent mode with the following steps: (1) full MS scan was acquired at 75000 of resolution with an AGC target to 3*106 in a maximum of 250ms for ma mass range of 350 to 1400 m/z; (2) MS/MS scan was acquired at 17500 of resolution with an AGC target of 1*105 in a maximum of 120ms and an isolation window of 1.5 m/z. Step 2 was repeated for the 8 most intense ions in the full scan (1) if the intensity was greater than 8.3*103 and the charge was 2 or 3. The peptide match option was set to on and isotopes of the same ion were excluded. Dynamic exclusion was set to 50s.

### 2.6 Data analyses and protein identification

Data files were converted to open source mzXML format using msConvert software from the ProteoWizard 3.0.9576 package (Kessner et al., 2008). During conversion, the MS and MS/MS data were centered. *Arabidopsis thaliana* protein database (Araport11) was used as a reference for protein identification. A contaminant database containing the sequences of standard contaminants such as trypsin, keratin, and serum albumin was also queried. Search was performed using X!Tandem (version Piledriver 2015.04.01.1) (Craig and Beavis, 2004). Tryspin was set in strict mode with 1 miscleavage in the first step. Carbamidomethylation of cysteine was set as a fixed modification. Oxidation of methionine, excision of N-terminal methionine, with or without acetylation, and pyroglutamate from glutamine or glutamic acid were set as potential modifications. In a second pass, the maximum allowed miscleavage was set to 5, deamidation on asparagine and glutamine and oxidation of tryptophan were added to the list of potential modifications.

Proteins were filtered and sorted using X!TandemPipeline (version 0.4.17) (Langella et al., 2017). Each identified protein was validated by the presence of at least two peptides with an E-value <0.01 and a protein E-value <10^-5^. According to these parameters, results were filtered to an estimated false discovery rate (FDR) of 0.15% at the peptide level and 0% at the protein level. The identified peptides were quantified by eXtracted Ion Current (XIC) and MassChroQ software (version 2.2.22) (Valot et al., 2011) using the following alignment parameters: ms2_1 alignment method tendency_halfwindow of 10, ms2_smoothing_halfwindow of 15, ms1_smoothing_halfwindow of 0. The quantification method XIC was based on max, min and max ppm range of 10, anti-spike half of 5, mean filter half_edge of 2, minmax_half_edge of 4 and maxmin_half_edge of 3. The thresholds for detection were 30,000 for min and 50,000 for max. The mass spectrometry proteomics data (**Table S1)** have been deposited to the ProteomeXchange consortium via the PRIDE partner repository with the dataset identifier PXD048867.

### 2.7 Peptide and protein normalization and quantification

The intensity of each peptide in each sample was normalized using a median-based method taking into consideration the peptide intensities of reference samples (Lyutvinskiy et al., 2013). The reference samples correspond to the pool of peptide ions extracted from the 18 samples analyzed using the same pipeline. Proteins were then quantified after removing common and doubtful peptide ions, peptides with too many missing data (more than 10%), peptides whose intensity profile deviated from the average profile of peptide-mz for the same protein, and the proteins quantified by a low number of peptide-mz (at least 2 peptides for a protein). Protein abundance was calculated as the sum of peptide intensities and log10 transformed (**Table S2)**.

### 2.8 Statistical, protein clustering and bioinformatics analyses of identified proteins

The results were analyzed using the program MCQR (version 0.5.2). Proteins regulated by U treatment were identified using a one-way analysis of variance (ANOVA) with the following linear model: Yij=µ + Ui + Ɛij, where Yij refers to individual protein abundance, *μ* is the general mean, Uj is the effect of U, and *ε*jk is the residual. For each protein, the P value obtained from the ANOVA test was adjusted (*Padj* < 0.05). Proteins that showed *Padj* <0.05 in the ANOVA were subjected to a Tukey test to determine the proteins that showed a difference between two doses (*P value* < 0.05) (**Table S3**). The Self-Organizing Tree Algorithm (SOTA) of the identified proteins was performed using the Z-score transformed values with eight clusters (**Table S4**). In **Table S5**, the AGI code, UniProt ID, gene name(s), gene description(s) for each identified protein were retrieved from the Araport 11 (Arabidopsis information resource (https://www.arabidopsis.org/) or UniProtKB databases. Protein classes were compiled from PantherDB (http://www.pantherdb.org/; last update February 23, 2022), whereas enzyme families were provided from Gene Ontology (GO) “molecular function” annotations available through QuickGO (www.ebi.ac.uk/QuickGO/) coupled with a manual curation (**Table S6**). The nature of (putative) metallic cofactors was obtained from the UniProtKB database (https://www.uniprot.org/). Predicted or experimentally proven subcellular localizations of proteins were compiled from the resource SUBAcon (SUBcellular Arabidopsis consensus v5; http://SUBA.live; last update June 30, 2022) (Hooper et al., 2017).

Hierarchical clustering was generated for the proteins that showed significant variation under U exposure using the maximum distance as the criterion dissimilarity. The Cytoscape v3.9.1 plugin Biological networks gene ontology (BiNGO) v3.0.3 (Maere et al., 2005) and the Metascape software (https://metascape.org) (Zhou et al., 2019) were used to calculate GO term enrichment of regulated proteins. The analysis was conducted using the default BINGO settings with the Bonferroni family-wise error rate correction (with significant level set at <0.05) and the *Arabidopsis thaliana* annotation. Bubble plots were generated with SR plot (https://www.bioinformatics.com.cn/en). For relevant positioning in metabolic pathways, proteins were analyzed using the Arabidopsis metabolic network knowledge database ChloroKB (Gloaguen et al., 2017).

## 3 RESULTS

### 3.1 Uranium is preferentially bound to membrane-associated and other insoluble proteins in roots

To gain insight into the dynamics of the root proteome under U stress, 5-week-old *Arabidopsis thaliana* (Col-0) plants were challenged with 5 and 50 µM uranyl nitrate for 48 h, thereafter referred to as U5 and U50 conditions, respectively. Uranyl nitrate was provided in distilled water in order to limit U interaction with other mineral components (*e.g.* phosphate) and maximize its absorption by roots (Misson et al., 2009; Doustaly et al., 2014; Sarthou et al, 2022). Roots treated with 5 and 50 µM uranyl nitrate accumulated 135 ± 23 and 1609 ± 157 µg U per g fresh weight, respectively (mean ± SD, n=6 biological replicates) (**Figure S1**). In shoots, U content was much lower (0.4 ± 0.1 and 2.2 ± 1.4 µg.g^-1^ FW in U5 and U50 plants, respectively), in agreement with the low translocation rate of U observed in Arabidopsis in these conditions (Vanhoudt et al., 2010; Laurette et al., 2012; Berthet et al, 2018; Sarthou et al, 2022). Total proteins were extracted from root tissues and soluble proteins were separated from insoluble material by ultracentrifugation. Then, membrane-bound and other insoluble proteins were extracted in the presence of 1% (w/v) SDS and recovered from residual insoluble material by centrifugation. Immunoblot analysis with antibodies against the membrane integral tonoplast protein TIP1;2 and the soluble fructose 1,6-bisphosphate aldolase isoforms (FBAs) was performed to assess the quality of protein fractions. TIP1;2 was detected only in the SDS-solubilized fraction, whereas aldolases were detected only in the soluble fraction (**Figures 1A, S2 and S3**). This result confirms the specific enrichment of the two fractions in either soluble or membrane-associated proteins with very low cross contamination. More details on the nature of the proteins enriched in the SDS-solubilized fraction will be provided by the proteomic analysis described below. Proteins from the soluble and SDS-solubilized fractions were mineralized and U was determined by ICP-MS. In the soluble fraction, the amount of U was 0.20 ± 0.05 and 3.02 ± 1.16 µg.mg^-1^ protein in the roots of U5 and U50 plants, respectively (**Figure 1B**). The amount of U recovered in membrane-associated and other insoluble proteins was about 10 to 15-time higher, with 3.1 ± 1.0 and 28.1 ± 3.6 µg U per mg protein (**Figure 1B**). This result is in agreement with previous data obtained in Arabidopsis cell suspension cultures treated with uranyl nitrate, in which most of U in the protoplasts was associated with membranes and only a small amount was found in the cell-soluble fraction (Sarthou et al., 2020). It also suggests that membrane-associated and other insoluble proteins are the preferred targets of U, and that their function and abundance may be particularly compromised under U stress. For these reasons, we focused our analysis on the dynamics of the root insoluble proteome in response to U stress.

**Figure 1.**
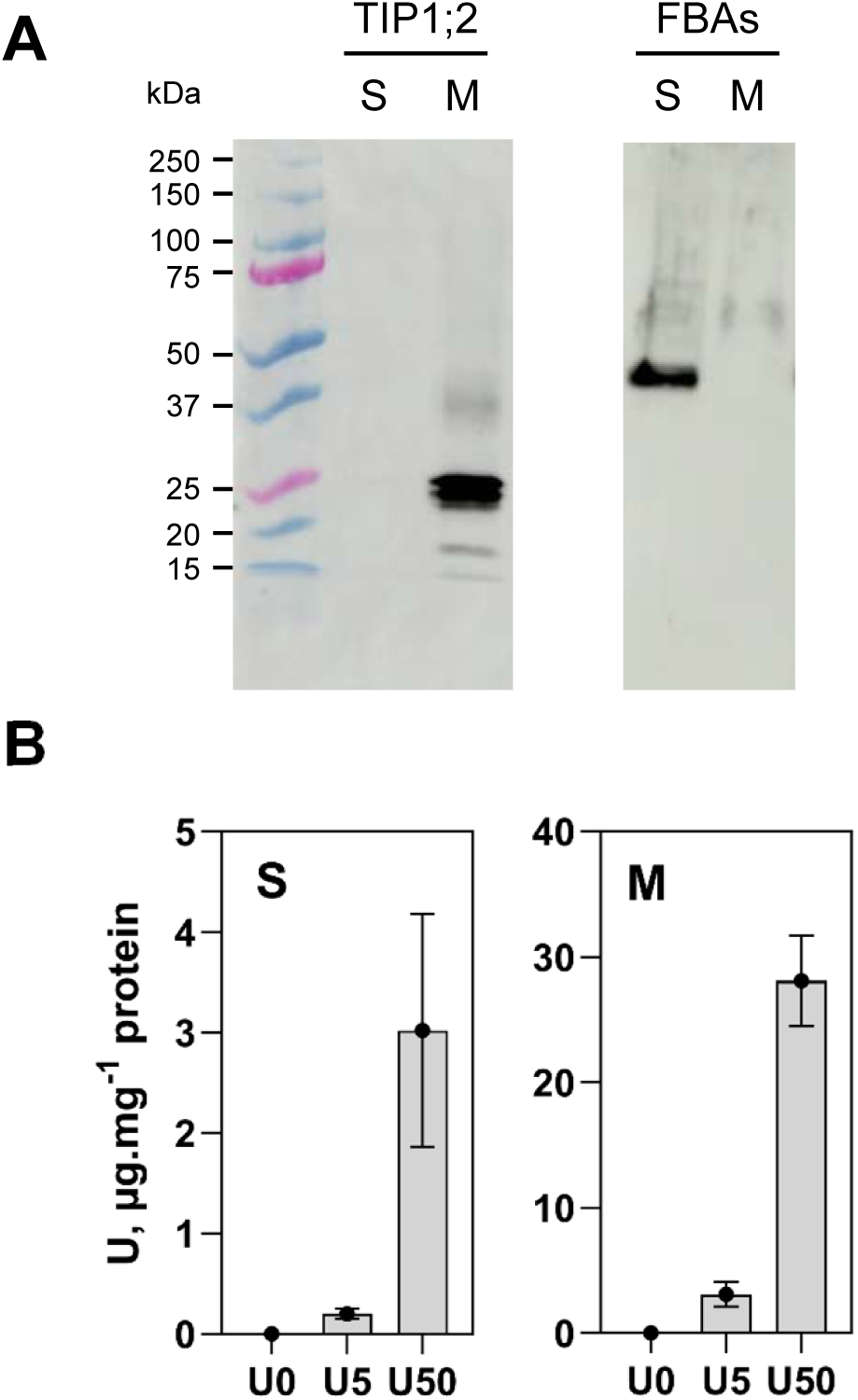
Quantification of uranium in soluble and membrane protein fractions from Arabidopsis roots. **(A)** Immunodetection of the tonoplast intrinsic protein TIP1;2, and plastid fructose-bisphosphate aldolases (FBAs) in soluble and membrane fractions isolated from Arabidopsis roots. One replicate of the experiment is show (untreated plants, samples R-0-2s and R-0-2m). SDS-PAGE and western blot analysis of all replicates are shown in Figures S2 and S3. (**B**) Uranium quantification in soluble and membrane protein fractions. Proteins were mineralized in nitric acid and U was quantified by ICP-MS. Bar plots represent mean ± SD with n=6 biological replicates.

### 3.2 The root membrane- and cell wall-enriched proteome is strongly affected by uranium

SDS-solubilized protein fractions from roots of the control (U0), U5 and U50 conditions were analyzed by nano-LC-MS/MS using a Q-Exactive mass spectrometer (n=6 biological replicates per condition). The analysis allowed the identification of 29,371 peptides corresponding to 3,462 indistinguishable protein groups (validated by at least 2 peptides with E-value<0.01, and a protein E-value<10^-5^) (**Table S1**). Whitin this dataset, 2,802 proteins (corresponding to 17,730 peptides) were quantified in a reproducible manner using a XIC-based approach (**Table S2**). The subcellular localization of the 2,802 proteins was analyzed using two complementary tools for gene ontology (GO) enrichment analysis, the BiNGO plugin of Cytoscape and the Metascape software for *A. thaliana*. The BiNGO analysis indicated that the most enriched (from 4- to 10-fold) subcellular compartments in the insoluble proteome are the peroxisome, plasma membrane, cell wall, endosome, nucleolus, vacuole, endoplasmic reticulum (ER), and nuclear envelope (**Figure S4A**). In addition, the Metascape cellular component enrichment analysis indicated that several protein complexes and vesicular systems were significantly enriched in this fraction (**Figure S4B**). Together, these results showed that the SDS-solubilized protein fraction from roots is enriched in membrane proteins from different subcellular compartments, in cell wall-associated proteins (enrichment >3.5), and in protein complexes (enrichment >3). This fraction is now referred to as the root membrane- and cell wall-enriched proteome, or simply the root membrane proteome.

Using a 1.5-fold change threshold for biological significance, we identified 458 differentially accumulated proteins (DAPs) in the root membrane proteome under U stress (ANOVA, p<0.05) (**Table S3**). As shown in Volcano plots (**Figure 2**), 68 proteins changed significantly between U0 and U5 (49 up-regulated, 19 down-regulated), 343 proteins changed between U0 and U50 (168 up-regulated, 175 down-regulated), and 299 proteins changed between U5 and U50 (143 up-regulated, 156 down-regulated). Overall, this indicated that most of the 458 DAPs were deregulated by a high U concentration (U50) whereas lower U stress (U5) moderately interfered with the root membrane- and cell wall-enriched proteome. To refine this comparison, hierarchical clustering analysis of the 458 DAPs clearly confirmed the grouping of biological replicates (n=6) for each U treatment (**Figure 3**). For each of the three conditions, a specific pattern of protein accumulation was observed. The root membrane proteome from plants cultivated in U0 and U5 conditions segregated drastically from that of U50. Finally, a self-organizing tree algorithm (SOTA) analysis was used to define eight clusters of proteins according to their accumulation patterns in relation to U concentrations in the medium (**Figure 4; Table S4**). Clusters 1 and 2 corresponded to proteins that preferentially accumulated as the U concentration increased. In contrast, clusters 4 and 5 identified proteins with decreasing abundance at increasing U concentrations. Clusters 3, 6, 7 and 8 grouped together proteins with opposite abundances at the two U concentrations. Approximately half of the 458 DAPs were up-regulated in response to U50 (clusters 1-3), whereas the abundance of the remaining 50% proteins was decreased in this condition (clusters 4-6) (**Figure 4**).

**Figure 2.**
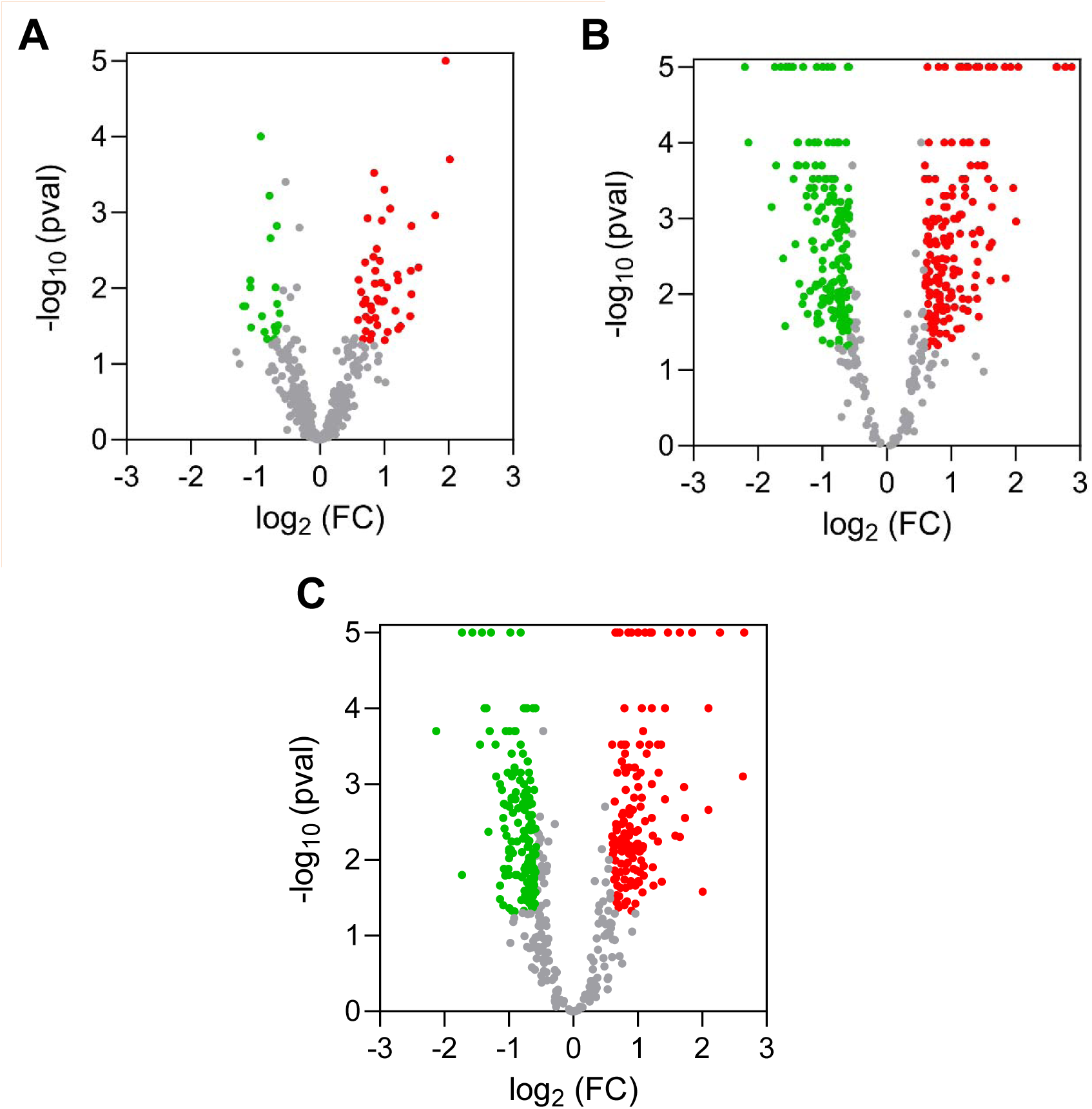
Volcano plots of root membrane proteins differentially accumulated during uranium stress. The relative abundance of root membrane proteins was compared in conditions U5 vs U0 (**A**), U50 vs U0 (**B**), and U50 vs U5 (**C**). Differentially accumulated proteins were defined using a fold change threshold >1.5 and a p-value <0.05 (Tukey test). Down-regulated and up-regulated proteins are shown in green and red, respectively. Proteins considered not regulated by U (fold change ≤1.5 and/or pvalue ≥0.05) are in grey. Proteins with p-value = 0 (Tukey test) were plotted with a –log10 (pval) of 5 for convenient graphical display.

**Figure 3.**
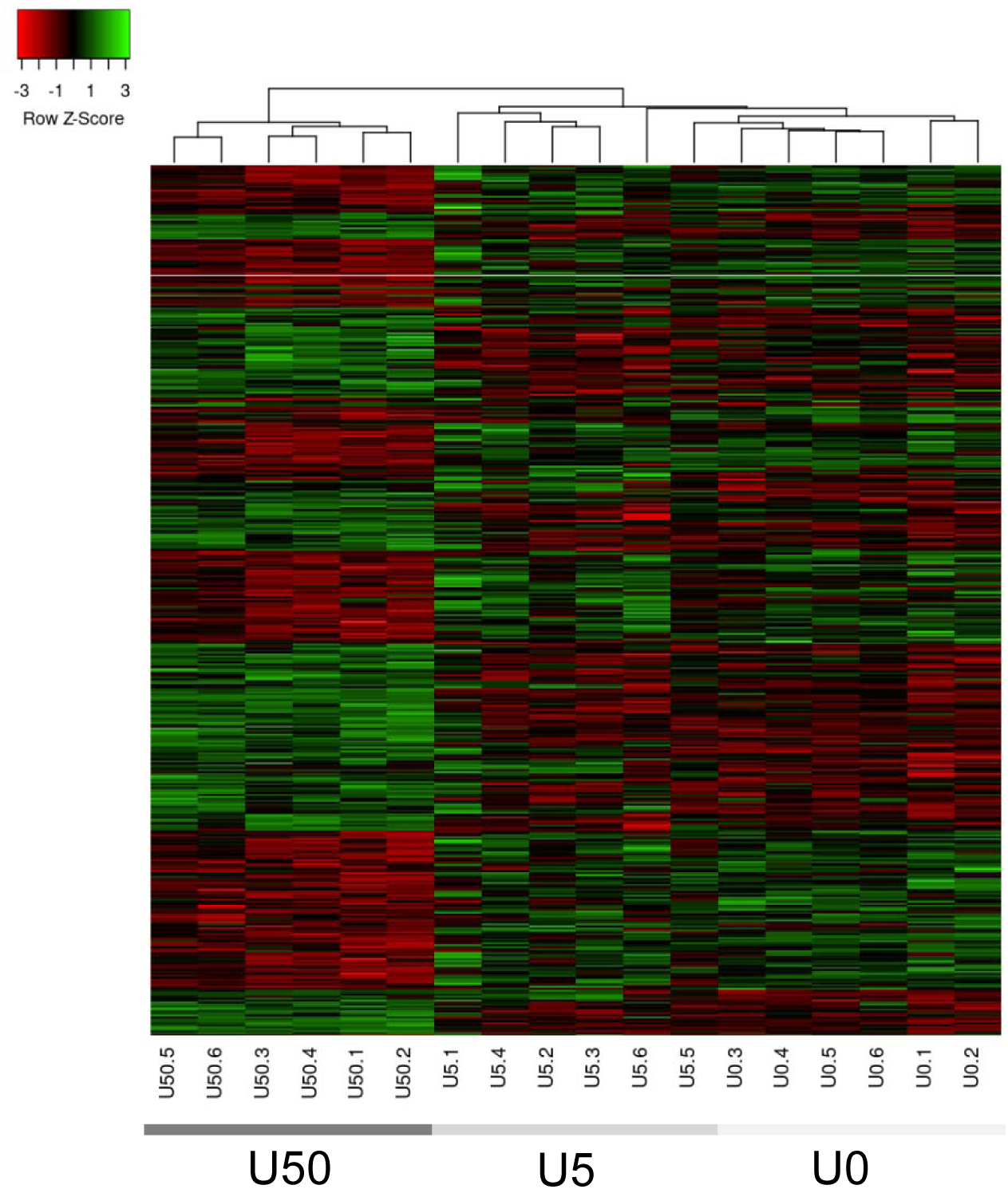
Global accumulation profiles of *A. thaliana* root membrane proteins in response to uranium stress. The heatmap represents the 458 proteins showing a significant change in abundance when exposed to 5 or 50 µM uranyl nitrate. Clustering was performed using the Heatmapper expression server with the average linkage clustering method and the Euclidean distance measurement method. Z-score normalization of protein expression values was done prior to clustering. The row Z-score is indicated.

**Figure 4.**
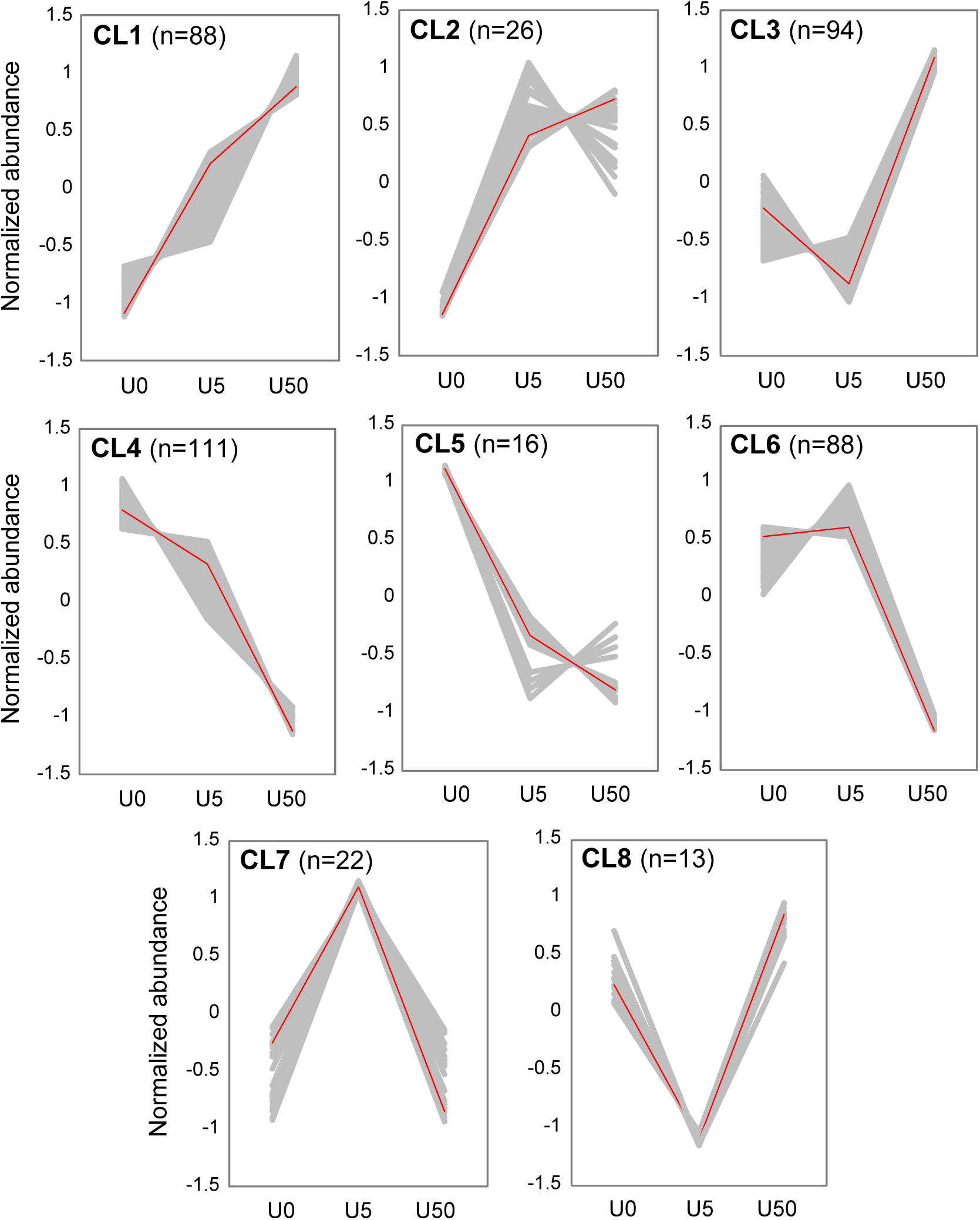
Clustering of proteins according to their accumulation patterns in response to uranium stress. Clustering was calculated by the SOTA method using Z-score transformed values to identify homogeneous patterns of protein abundance changes. Eight clusters (CL1 to CL8) have been defined. Individual profiles are depicted by gray lines (Z-score), the average profile is marked in red. The number of proteins in each cluster is indicated.

### 3.3 Membrane- and cell wall-associated proteins regulated by uranium perform a wide range of molecular functions

To further delineate whether some cell compartments are specifically impacted by U stress, the subcellular localization of the 458 DAPs was analyzed using the BiNGO and Metascape tools. The plasma membrane, ER, vacuole, and cell wall were the most enriched subcellular compartments in the BiNGO analysis (>3.5-fold enrichment) (**Figure 5A**). Additionally, the supramolecular complex, secretory vesicle, Golgi cisterna, and Casparian strip terms were overrepresented in the Metascape analysis (>3- fold enrichment) (**Figure 5B**). Altogether, these results suggest that U influences the abundance of proteins distributed in most of the cell compartments, and support an enrichment of the cell wall and membrane fractions. A moderate enrichment (2.7-fold) of proteins detected in the cytosol could reflect the presence of protein complexes interacting, at least transiently, with the membranes.

**Figure 5.**
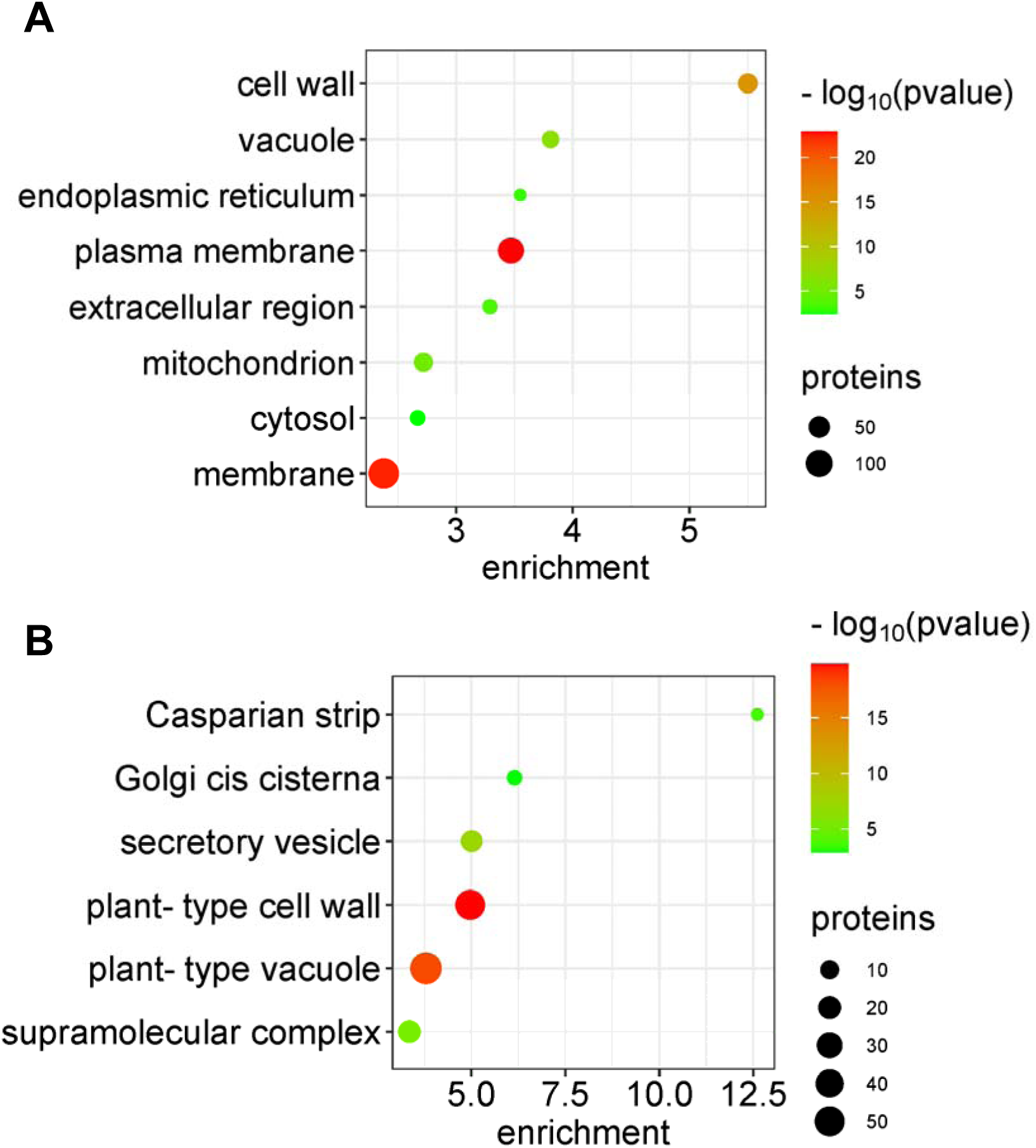
GO enrichment analysis of cellular components regulated by uranium stress. GO enrichment analysis of cellular components was performed with the 458 DAPs using the (**A**) BiNGO and (**B**) Metascape tools. Bubble plots show GO terms ordered by enrichment factors (threshold >2.3 for BiNGO, >3 for Metascape). BiNGO settings to assess overrepresented GO cellular components were as follows: statistical hypergeometric test, Bonferroni Family-Wise Error rate multiple testing correction, and significant p-value <0.05. Genes were annotated with plant GO slim terms. The Metascape enrichment analysis has been done with the GO cellular components ontology source. Terms with a p-value <0.01, a minimum count of 3, and an enrichment factor >1.5 have been grouped into clusters based on their membership similarities.

We investigated the predicted molecular functions of the 458 DAPs under U stress. Proteins were grouped into functional classes based on GO annotations related to the molecular function and data were manually curated. Our analysis showed that enzymes represented 46% of U-regulated proteins (**Figure 6A**; **Table S6**). The other proteins belonged to the classes of transporters (13%), transcriptional/translational factors (9%), chaperones/co-chaperones (6%) or transmembrane receptors (4%), while 24 proteins (5%) had other functions and 89 proteins (19%) were referred to as “unknown” (**Figure 6A; Table S5**). Focusing on the enzyme class, hydrolases, oxidoreductases and transferases were mainly identified, while lyases, ligases and isomerases accounted for a smaller proportion (**Figure 6A; Table S5**). The hydrolase protein class included proteases, phosphatases, glycosidases, esterases, lipases, nucleases and helicases (**Table S5**). Oxidoreductases were primarily represented by peroxidases. Finally, kinases, transaminases and aldolases formed the transferase class. Concerning transporters, families involved in the transport of nucleotides, proteins, vesicles, water, sugars and metals were regulated by U (**Table S5**). Of note, more than 30% of the proteins identified during our proteomic analysis were annotated as metallic proteins (**Figure 6B**). The abundance of genuine or predicted iron-, calcium-, zinc-, and magnesium-containing proteins was peculiarly affected by U (**Figure 6B**).

**Figure 6.**
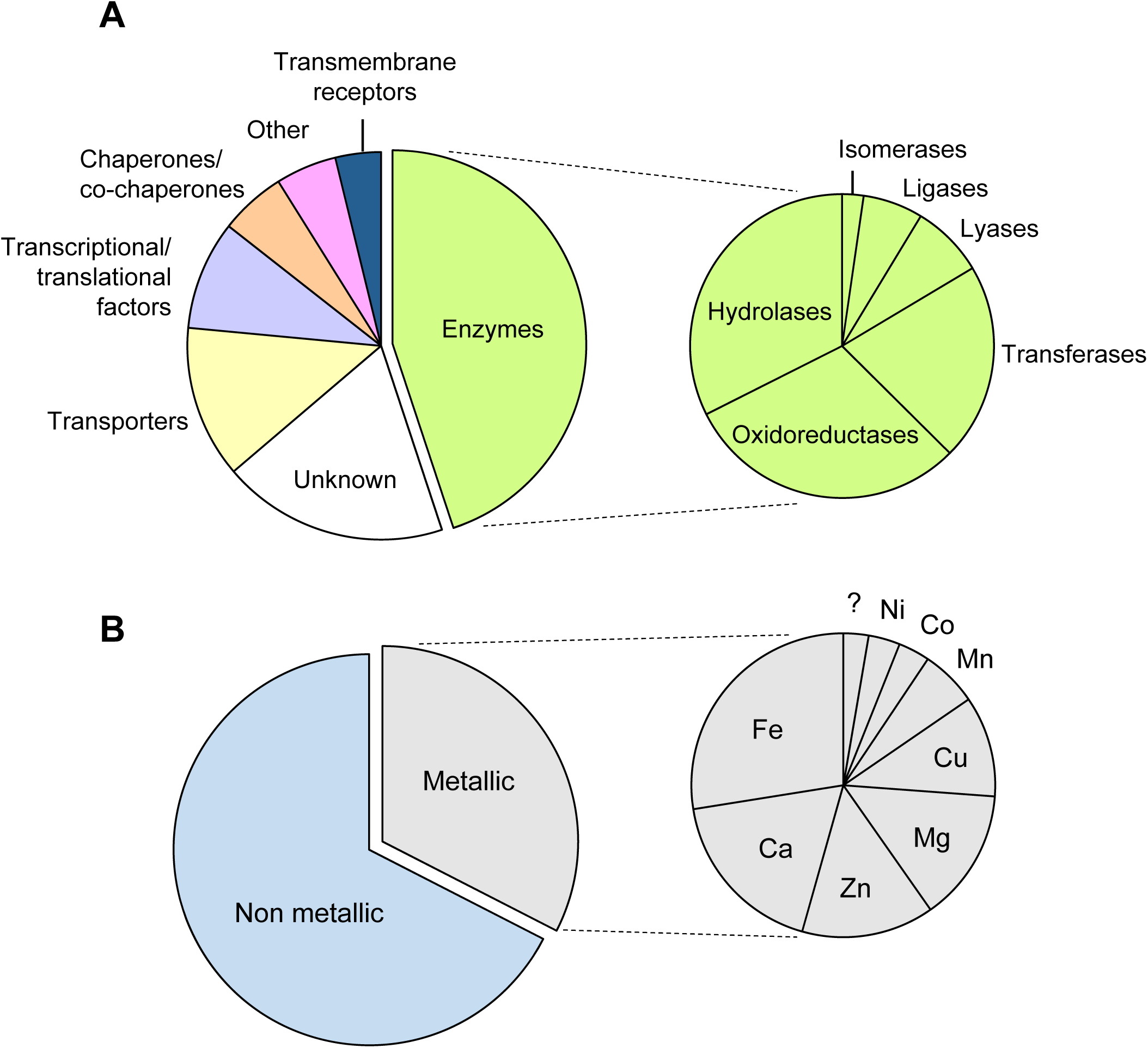
Protein classes and metal cofactors associated with differentially accumulated proteins under uranium stress. Circular diagrams representing (**A**) the proportion of the different protein classes and (**B**) the nature of cofactors associated with the identified proteins. Fe: iron, Ca: calcium, Zn: zinc, Mg: magnesium, Cu: copper, Mn: manganese, Co: cobalt, Ni: nickel, ?: unknown metal.

### 3.4 Multiple biological pathways involved in stress response, metabolism and cell organization are modified by uranium

To provide an overview of the biological processes regulated by U, we performed an enrichment analysis of the 458 DAPs using BiNGO and Metascape. For biological process enrichment analysis, the Metascape tool has the advantage of identifying GO constituted by a smaller set of proteins compared to BiNGO. Among the most enriched (>1.8-fold) biological processes, and considering only higher levels in the GO term hierarchy, the terms response to stress (GO:0006950), response to metal ion (GO:0010038), amino acid metabolic process (GO:0006520), monocarboxylic acid metabolic process (GO:0032787), and transport (GO:0006810) were identified using BiNGO (**Figure 7A**). The analysis using Metascape revealed additional enriched (>2.2-fold) biological processes such as protein folding (GO:0006457), tRNA aminoacetylation (GO:0043039), phenylpropanoid metabolic process (GO:0009698), cell wall organization (GO:0071555), and cell-cell junction assembly (GO:0007043) (**Figure 7B**). This analysis indicates that multiple biological processes are affected in response to U. In the following paragraphs, we describe in more detail the molecular actors contained in GO terms and sub-terms. To provide a clear overview of the effects of U on the root membrane proteome, we have grouped these terms/sub-terms into three main categories, namely (i) stress, (ii) metabolism and cell wall organization, and (iii) transport and compartmentalization.

**Figure 7.**
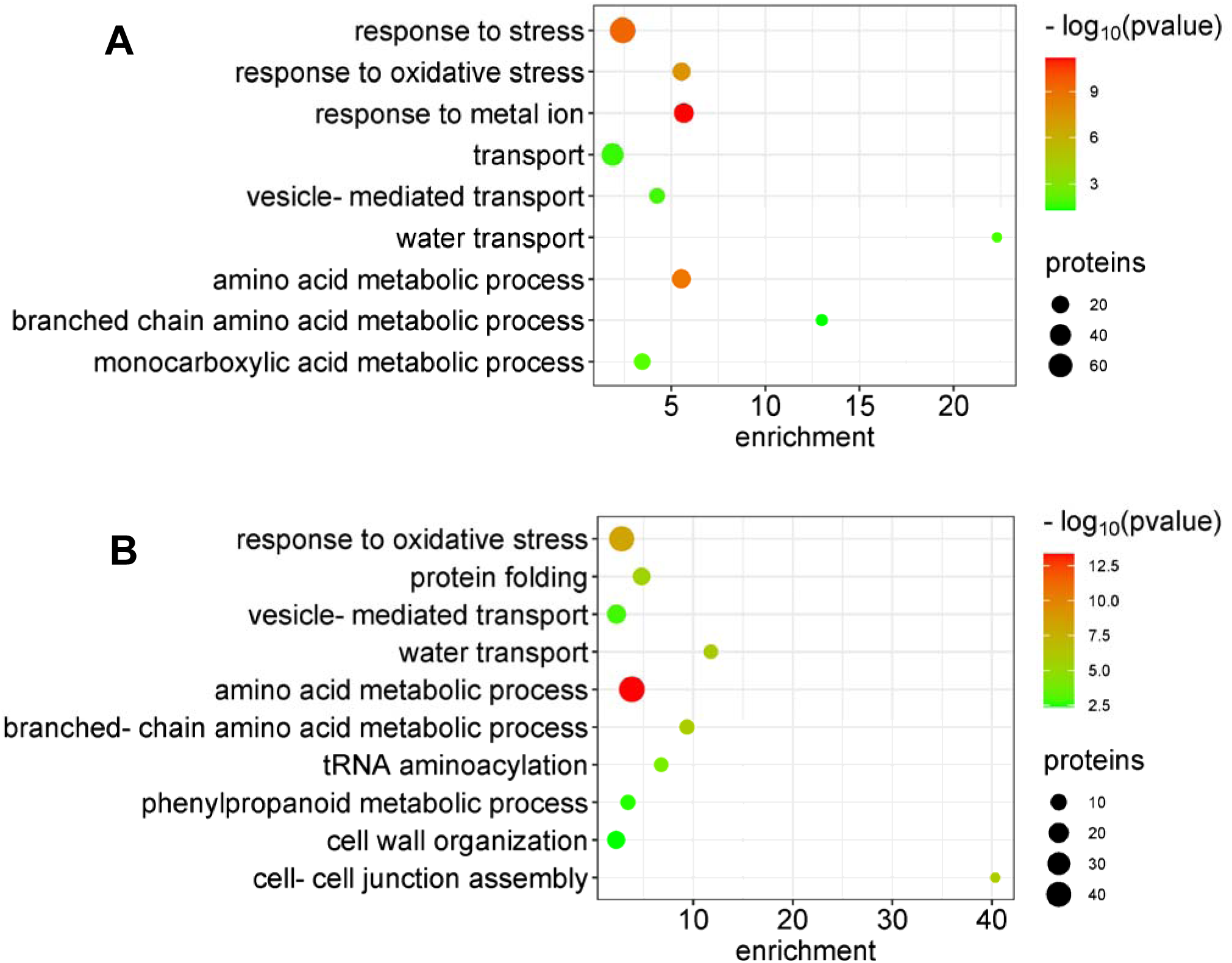
GO enrichment analysis of biological processes regulated by uranium stress. GO enrichment analysis of biological processes was performed with the 458 DAPs using the (**A**) BiNGO and (**B**) Metascape tools. Bubble plots show GO terms grouped by major functions (threshold >1.8 for BiNGO, >2.2 for Metascape). BiNGO settings to assess overrepresented biological processes were statistical hypergeometric test, Bonferroni Family-Wise Error rate multiple testing correction, and significant p-value <0.05. The Metascape enrichment analysis has been done with the GO biological processes ontology source. Terms with a p-value <0.01, a minimum count of 3, and an enrichment factor >1.5 have been grouped into clusters based on their membership similarities.

#### Stress

The stress category includes the GO terms response to stress (and its child term response to oxidative stress, GO:0006979), protein folding, and response to metal ions. Response to stress is one of the largest GO biological process with 75 DAPs mainly present in clusters CL1, CL3, CL4 and CL6 (**Figure 4**). Among them, 23 proteins constitute the sub-group response to oxidative stress (**Figure 8A**). Only a few of these proteins were accumulated upon U stress (CL1 and CL3), including plastidial peroxiredoxin PRXIIB and PRXIIE, mitochondrial peroxiredoxin PRXIIF, glutathione peroxidase 6 (GPX6), thioredoxin H type 5 (TRX5), and peptide methionine sulfoxide reductase 4 (PMSR4). All other proteins in this group were less abundant in the presence of U50, such as cytosolic and plastidial copper/zinc superoxide dismutase 1 and 2 (CSD1/2), gamma-glutamyl transpeptidase 1 (GGT1), and up to 13 class-III peroxidases (PRXs) (**Figure 8A**). The second GO sub-group, protein folding, comprised a dozen proteins accumulating in response to U (CL1 and CL3). These included, in particular, several chaperones/chaperonins, and three peptidyl-prolyl cis-trans isomerases (ROC1/3/5). Finally, many of the DAPs involved in response to stress (at least 32 proteins) are metalloproteins and/or proteins regulated by metal ions. In addition to the numerous PRXs and CSDs already described above, this group contains a number of proteins involved in the response to biotic and abiotic stress and/or metal homeostasis. Among the up-regulated proteins, the calcium- and zinc-binding chaperone calreticulin-3 (CALR3), the calcium-, zinc- and iron-binding early response to dehydration 10 and 14 proteins (ERD10/14), the nickel-binding phosphorylated protein 34 (PHOS34), as well as the cadmium- and zinc-responsive pre-mRNA regulatory protein glycine-rich protein 7 (GRP7) were found. In contrast, the chloroplastic iron storage protein ferritin 1 (FER1) and the plasma membrane-associated cation-binding protein 1 (PCaP1) were present at lower levels under U stress.

**Figure 8.**
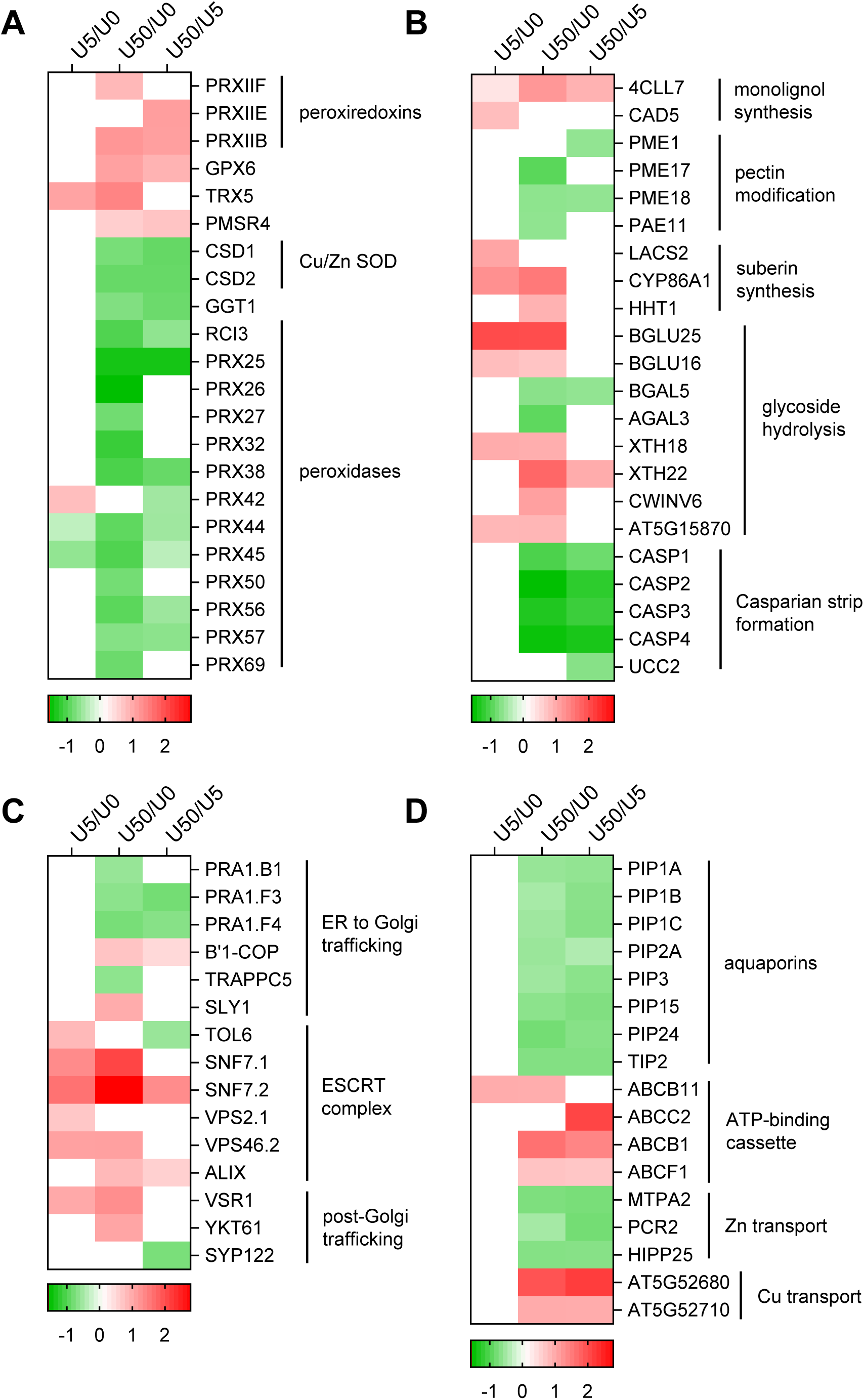
Heatmaps representing sets of proteins regulated by uranium stress. Proteins whose abundance is significantly (p<0.05) increased or decreased in response to U stress are shown in red and green, respectively. Protein changes not supported by p<0.05 are in white. Expression levels values are in log2 scale. (**A**) stress, (**B**) cell wall organization, (**C**) vesicular transport, (**D**) solute transport.

#### Metabolism and cell wall organization

Several terms related to plant metabolism and cellular organization are significantly enriched among the 458 DAPs. They include amino acid metabolic process (and its sub-term branched chain amino acid metabolic process, GO:0009081), monocarboxylic acid metabolic process, tRNA aminoacetylation, phenylpropanoid metabolic process, cell wall organization, and cell-cell junction assembly (**Figure 7**). These proteins are distributed in all eight clusters from the SOTA analysis, but are mainly present in CL1, CL3, CL4, and CL6. The GO term amino acid metabolic process includes enzymes involved in the synthesis of several amino acids (**Figure S5**). The synthesis of branched-chain amino acids (BCAA; Ile, Leu, Val) is particularly impacted by U stress with the accumulation of four enzymes catalyzing consecutive steps, namely acetohydroxy acid synthase (ALS), acetohydroxy acid isomeroreductase (ILV5), dihydroxyacid dehydrate (DHAD), and isopropylmalate synthase (MAML-4) (**Figure S5**). In contrast, the isopropylmalate isomerase 2 (IPMI2), methylthioalkylmalate synthase 3 (MAM3), superroot (SUR1) and cytochrome 5B-C proteins involved in the biosynthetic pathway of BCAA-derived aliphatic glucosinolates are less abundant (Harun et al., 2020). The synthesis of aromatic amino acids (Phe, Tyr, Trp), aspartate-derived amino acids (Thr, Lys), amino acids contributing to one-carbon metabolism (Ser, Gly, Met), and Arg are also up-regulated in response to U50 (**Figure S5**). The highest concentration of U also has a significant effect on tRNA aminoacetylation reactions as observed by the accumulation of eight aminoacyl tRNA-ligases (**Figure S5**), suggesting an effect of the radionuclide on *de novo* protein synthesis. In connection with aromatic amino acid synthesis, the response to U stress is characterized by the differential accumulation of enzymes belonging to the phenylpropanoid metabolic process. Two enzymes involved in the activation of phenylpropanoid precursors, coumarate CoA ligase (4CLL7) and cinnamyl alcohol dehydrogenase (CAD5), accumulated during U stress (**Figure 8B**), suggesting an increased synthesis of flavonoids or monolignols (Fraser and Chapler, 2011). Following their transport to the cell wall, monolignols are oxidized by members of the large family of class-III peroxidases and further polymerized to form lignin. As mentioned before, 13 class-III peroxidases predicted to be extracellular were decreased in abundance in response to U exposure (**Figure 8A**), suggesting a significant effect of U on the homeostasis of this major cell wall polymer. In addition, important changes in cell wall composition are supported by several proteins whose abundance changes during U stress. First, the abundance of enzymes involved in the modification of pectin, namely pectin methylesterases (*i.e.* PME1/17/18) and pectin acetylesterase 11 (PAE11), is decreased in response to U stress (**Figure 8B**). Second, enzymes involved in either suberin synthesis (*e.g.* long chain acyl-CoA synthetase LACS2, cytochrome P450 86A1, feruloyl-CoA transferase HHT1) (Serra and Geldner, 2022) or the degradation of cell wall polysaccharides (glycoside hydrolases acting on diverse substrates, *i.e.* alpha and beta-galactoside, beta-glucoside, xyloglucan, fructan) (Minic and Jouanin, 2006) are either more or less abundant in U5 and U50 than in control samples (**Figure 8B**). These important changes in cell wall-organizing enzymes suggest a dynamic and complex reorganization of the cell wall in response to U stress. This assumption is strengthened by the observation that the abundance of four Casparian strip membrane proteins (*i.e.* CASP1-4) and uclacyanin 2 (UCC2), both involved in the Casparian strip formation and consequently in cell-cell junctions (Roppolo et al., 2011; Reyt et al., 2020; Barbosa et al., 2023), was decreased in the presence of U50 (CL4 in **Figure 4**, **Figure 8B**).

#### Transport and compartmentalization

The transport group, formed by the enriched GO term transport and its child terms vesicle-mediated transport (GO:0016192) and water transport (GO:0006833), contains 47 DAPs associated with CL1, CL2 and CL6 (**Figure 4**). The child terms vesicle-mediated transport and water transport were enriched up to 4- and 22-fold, respectively, according to the BiNGO and Metascape analyses (**Figure 7**). An important regulation of the endomembrane trafficking pathways in response to U was suggested by the differential accumulation of the prenylated rab acceptors PRA1 (*i.e.* PRA1.B1/F3/F4), the β’1 subunit of the COP1 coat, the transport protein particle C5 (TRAPPC5) subunit of the TRAPP I complex, and the Sec1/Munc18 protein SLY1 (**Figure 8C; Figure S6**). All these proteins mediate the vesicle transport between the ER and the Golgi apparatus. Acting downstream, the ESCRT-0 like TOM1-like protein 6 (TOL6) and some ESCRT-III-related components, including the sucrose non-fermenting 7.1/7.2 (SNF7.1/7.2), vacuolar protein sorting 2.1/46 (VPS2.1/46), and alg-2 interacting protein-x (ALIX) were accumulated upon U stress. Only the abundance of TOL6 was reduced at U50. These sequential ESCRT complexes orchestrate the biogenesis of multivesicular body (MVB) and the sorting of ubiquitinated cargo proteins for vacuolar degradation (Gao et al., 2017). The abundance of other proteins involved in the post-Golgi trafficking were accumulated during U stress, including the vacuolar-sorting receptor 1 (VSR1), and the SNARE proteins YKT61 and SYP122 (**Figure 8C; Figure S6**). Concerning water transport, eight aquaporins belonging to the plasma membrane intrinsic protein (PIP) and tonoplast intrinsic protein (TIP) subfamilies were down-regulated in the presence of a high concentration of U (**Figure 8D**). Finally, among transporters likely to transport metals, the zinc transporters MTPA2, PCR2 and HIPP25 were down-regulated under U50 whereas the At5g52680 and At5g52710 proteins annotated as copper transporters were up-regulated under these conditions (**Figure 8D**). The ABCB1, ABCB11, ABCC2, and ABCF1 members of the ABC transporter family, related to metal homeostasis, were specifically accumulated under different U concentrations (**Figure 8D**).

## 4 DISCUSSION

XIC-based quantification of the membrane- and cell wall-enriched proteome of Arabidopsis roots identified 2,802 proteins and showed that the abundance of 458 of them was significantly changed during U stress (fold change >1.5, p <0.05). The two high-affinity uranyl-binding proteins identified so far in Arabidopsis, PCaP1 and GRP7 (Vallet et al., 2023; unpublished data), are present among the 458 DAPs. This finding supports the view that the insoluble fraction of the root proteome, as a hot spot of U accumulation (**Figure 1**), is a relevant compartment to analyze the consequences of U intoxication and to reveal new defense mechanisms. About 30% of the identified DAPs are genuine or putative metalloproteins (**Figure 6**). This figure per se does not reflect an enrichment in metal-containing proteins as a commonly cited approximation is that one-third of proteins require metals, of which magnesium is the most common (Waldron et al., 2009). Yet, our analysis indicates that the abundance of iron- and calcium- containing proteins is the most affected by U (**Figure 6**). This observation corroborates previous data showing a preferential interference between U and iron or calcium homeostasis in plants (Doustaly et al., 2014; Berthet al., 2018; Mertens et al., 2022; Sarthou et al., 2022). One of the explanation would be that uranyl ions compete and displace iron and calcium in some proteins, as observed in the eukaryotic transferrin and calmodulin proteins (Vidaud et al., 2007; Pardoux et al., 2012). The release of free iron induced by U would lead to oxidative stress. In line with this hypothesis, a set of antioxidant enzymes and chaperones related to oxidative stress was significantly modified upon U stress (**Figure 8A**). An increase of ROS species has already been observed in roots from plants challenged with U (Aranjuelo et al., 2014; Tewari et al., 2015, Serre et al., 2019; Gupta et al., 2020; Xiao et al., 2023). Depending on the study, the activity of antioxidant enzymes were found up- or down- regulated. Our proteomic analysis showed that the abundance of antioxidant enzymes such as peroxiredoxins or superoxide dismutases (**Figure 8A**) was either increased or decreased in U-stressed Arabidopsis, indicating a coordinated and dynamic action of these enzymes to prevent or maintain ROS at low levels.

Our analysis showed an important deregulation of amino acid metabolism in response to U stress (**Figure 7; Figure S5**). However, it should be borne in mind that analysis of the root membrane- and cell wall-enriched proteome provides a partial view of amino acid metabolism, which is mainly soluble but involves multienzyme complexes (e.g. metabolons) or membrane-associated proteins (Zhang and Fernie, 2021). An interference between nitrogen metabolism and U has been previously described in plants and a metabolomic analysis of *Vicia faba* roots showed that U significantly reduced the content of various free amino acids (Lai et al., 2020; Chen et al., 2023). Our proteomic analysis revealed that several enzymes involved in the synthesis of various amino acids, in particular branched-chain amino acids (BCAA), were more abundant when U was applied (**Figure S5**). An increase in BCAA biosynthesis could be explained by their important role in plant responses to a wide range of abiotic stresses (Obata and Fernie, 2012), including cadmium stress (Zemanova et al., 2017; Zhao et al., 2020). In this context, the accumulation of Leu, Ile and Val may serve to promote stress-induced protein synthesis (Nambara et al., 1998). The accumulation of several aminoacyl tRNA ligases in response to U stress supports this hypothesis (**Figure S5**). The accumulation of enzymes involved in BCAA biosynthesis may also reflect a compensatory mechanism due to their rapid catabolism to cope with U stress. In fact, the breakdown products of BCAAs (*e.g.* acetyl-CoA, propionyl-CoA, acetoacetate) are potential energy sources for plants (Joshi et al., 2010). Another hypothesis would be that the synthesis of aliphatic glucosinolates derived from BCAAs may have a role in U detoxification and tolerance in Arabidopsis. In fact, four enzymes involved in the biosynthetic pathway of these specialized metabolites are down-regulated under high U concentration. A decrease in glucosinolates has been reported in Brassicaceae plants exposed to a cadmium stress (Sun et al., 2009; Jakovljevic et al., 2013). One hypothesis would be that plants limit channeling of organic sulfur into the glucosinolate pool in order to support the sulfate demand for glutathione and phytochelatin biosynthesis, which are important metabolites to limit heavy metal-related damages (Herbette et al., 2006). Besides, the biosynthetic pathway of aromatic amino acids was also increased in response to U stress (**Figure S5**). In this case, Trp could serve for the biosynthesis of phytohormones (*i.e*. auxin) whereas Phe could be required for the phenylpropanoid pathway to produce a wide range of plant secondary products, especially antioxidative metabolites (flavonoids, anthocyanins, lignins) and phenolic compounds in response to abiotic stress such as U.

Previous studies have shown that a high proportion of uranyl ions in plant roots are associated with the cell wall, which is rich in negatively-charged groups with high affinity for metal cations (Lai et al., 2020; Sarthou et al., 2020; Chen et al., 2023). Our analysis supports important changes in cell wall composition in response to U stress, including lignin and suberin synthesis, pectin modifications, polysaccharide hydrolysis, and Casparian strips formation (**Figure 8B**). The abundance of three pectin methylesterases and one acetylesterase was decreased during U stress (**Figure 8B**). A negative relationship between the degree of methylesterification and acetylation of pectins and their ability to bind some heavy metals (*i.e.* aluminum, lead, copper) has been reported in plants (Khotimchenko et al., 2007; Mimmo et al., 2009; Colzi et al., 2012). By maintaining a high degree of pectin esterification, plants could improve their tolerance to U by limiting heavy metal accumulation in their tissues, thus preventing transport to intracellular compartments. Another adaptive strategy in response to heavy metal intoxication in plants is cell wall thickening. The deposition of callose and lignin in lateral roots together with structural damage to root epidermal cells have been observed in response to U (Serre et al., 2019; Lai et al., 2020). Counter-intuitively, our analysis shows that the CASP protein family and UCC2, which are involved in the formation of lignin-based Casparian strips in the root endodermis (Roppolo et al., 2011; Reyt et al., 2020; Barbosa et al., 2023), are down-regulated during U stress (**Figure 8B**). Thus, a strong defect in root apoplastic permeability caused by a disruption of Casparian strips would facilitate the radial transfer of U in roots and ultimately its translocation toward aboveground organs. This is obviously not the case, as we observed a very low accumulation of U in the shoots of treated plants (**Figure S1**). Instead, the limitation in U translocation from roots to shoots would rather induce a U-dependent ectopic callose deposition accompanied by an enhanced suberization caused by a reduction in the abundance of CASP and UCC2 proteins, similar to what is observed in corresponding Arabidopsis loss-of-function mutants (Reyt et al., 2020; Barbosa et al., 2023). Several enzymes involved in suberin synthesis are up-regulated during U stress (**Figure 8B**), reinforcing this hypothesis.

A direct consequence of disrupted radial apoplastic root transport caused by ectopic lignin and suberin deposition could be the reduced abundance of eight aquaporins (**Figure 8D**) and the likely limitation of water flux in roots. Another explanation would be that changes in aquaporin levels is a preventive mechanism to limit the harmful effects of U in leaves by reducing water flow in roots. Indeed, water transport has been shown an important process regulating the partitioning and accumulation of U (Aranjuelo et al., 2014). Moreover, a permeability of aquaporins to metalloids has been reported in plants. This is particularly the case for NIP1.1, which has been shown to be involved in antimony and arsenite transport in Arabidopsis (Kamiya and Fujiwara, 2009; Kamiya et al., 2009). These observations suggest that disruption of specific aquaporins could confer U tolerance to plants.

Concerning U trafficking in plants, only the contribution of the calcium channels ANN1 and MCA1 to root U uptake has been experimentally demonstrated (Sarthou et al., 2022). Then, the identification of transporters involved in intracellular U trafficking remains to be deciphered, and only a few clues are available from transcriptomic data obtained from plants challenged with U (Doustaly et al., 2014, Mertens et al., 2020; Lai et al., 2020). Our proteomic approach points out the accumulation of several transporters belonging to the ABC transporter family, which has been shown to participate in heavy metal tolerance in plants (**Figure 8D**). The phytochelatin transporter ABCC2 may be of particular interest for U detoxification, as this protein confers tolerance to several heavy metals, including arsenic, cadmium, and mercury (Song et al., 2010; Park et al., 2012). Zinc, copper, cadmium or uncharacterized transport proteins are also deregulated by high U concentrations (**Figure 8D**). Together with ABC transporters, these proteins may represent relevant molecular actors in U tolerance, either by transporting the radionuclide or by maintaining the homeostasis of essential metals.

Finally, our analysis revealed the importance of endomembrane trafficking in the response of plants to U stress. Indeed, the abundance of many proteins involved in plant endomembrane trafficking was changed upon U stress (**Figure 8C; Figure S6**). More specifically, proteins involved in the traffic between the ER and Golgi apparatus, in pre-vacuolar compartments/multi vesicular body formation (*i.e.* ESCRT complexes), and in the secretory pathway were significantly deregulated (**Figure 8C; Figure S6**). The involvement of endocytosis in U uptake has been recently proposed in tobacco cells (John et al., 2022). Here, a hypothetical scenario would be that uranyl ions, U target proteins or other proteins involved in U tolerance are internalized into vesicles to prevent any toxicity before being released towards the apoplast or stored/degraded in the vacuole. As an example, the ESCRT-III accessory component ALIX was previously demonstrated to mediate sorting and vacuolar degradation of the high-affinity phosphate transporter PHT1;1 (Cardona-López et al., 2015). Since U forms chelates with phosphate, the observed accumulation of the ALIX protein in U-treated plants may represent a strategy to mitigate U toxicity.

To conclude, our results show that Arabidopsis roots orchestrate an important rearrangement of the cell wall and membrane proteome in response to U stress. Our proteomic data shed light on biological processes disrupted by U and improve our understanding of the mechanisms by which plants cope with metal toxicity. This study also provides a set of transporters and metal-binding proteins that may be involved in the fate of U in plants. Further functional studies will be required to elucidate the role of these proteins in U tolerance.

## 5 ACKNOWLEDGMENTS

We thank Célia Baggio (LPCV) for technical assistance with plant growth. We thank Dr Michel Zivy (PAPPSO) for the fruitful discussions in setting up the project. This work was funded by the Agence Nationale de la Recherche (ANR-17-CE34-0007, GreenU and ANR-17-EURE-0003, CBH-EUR-GS).

## 8 SUPPLEMENTARY MATERIAL

**Figure S1. Uranium content in roots and shoots of Arabidopsis plants.**

Uranium was measured by ICP-MS in roots (A) and shoots (B) of control (U0) and U-treated plants (U5 and U50 for 5 and 50 µM uranyl nitrate, respectively). Data are mean ± SD of n=6 biological replicates.

**Figure S2. SDS-PAGE analysis of soluble and membrane proteins from Arabidopsis roots.**

SDS-PAGE analysis of (**A**) soluble and (**B**) membrane proteins from control and U-treated *A. thaliana* roots. Proteins were stained with Coomassie Blue. Sample nomenclature: R, root; U concentration (0, 5, 50 µM uranyle nitrate): 1s to 6s, biological replicates of soluble protein extracts; m1 to m6, biological replicates of membrane protein extracts; *, samples analyzed by western blot (Figure S3).

**Figure S3. Quality assessment of membrane proteins from Arabidopsis roots by Western blot analysis.**

Western blot detection of (**A**) the fructose-bisphosphate aldolases (FBAs) and (**B**) the tonoplastic intrinsic protein 1;2 (TIP1;2) in soluble (s) and membrane (m) protein extracts isolated from roots of Arabidopsis plants treated with 0, 5 and 50 µM uranyl nitrate. SDS-PAGE analysis of protein extracts (including sample nomenclature) is shown in Figure S2.

**Figure S4. GO enrichment analysis of cellular components in the membrane and cell wall proteome of Arabidopsis roots.**

GO enrichment analysis of cellular components was performed using the 2,802 proteins identified by mass spectrometry using the BiNGO (**A**) and Metascape (**B**) tools. Bubble plots show GO terms ordered by enrichment values (threshold >4 for BiNGO, >2.5 for Metascape). BiNGO settings to assess overrepresented GO cellular components were as follows: statistical hypergeometric test, Bonferroni Family-Wise Error rate multiple testing correction, and significant p-value <0.05. The Metascape enrichment analysis has been done with the GO cellular components ontology source. Terms with a p-value <0.01, a minimum count of 3, and an enrichment factor >2.0 have been grouped into clusters based on their membership similarities.

**Figure S5. Effect of uranium on amino acid metabolism.**

(A) Proteins regulated by U are mapped to the KEGG pathway ‘biosynthesis of amino acids’ (ath01230). (B) Protein expression profiles are shown on a heatmap. The numbers in brackets refer to the enzyme positions in the pathway. (**C**) Heatmap of tRNA ligases regulated during U stress (not indicated on the pathway). Proteins whose abundance is significantly (p<0.05) increased or decreased in response to U stress are shown in red and green, respectively (log2 scale).

**Figure S6. Proteins differentially regulated by uranium involved in endomembrane trafficking.**

PRA1s play a role in the trafficking of cargo proteins destined to various endomembrane compartments (Jung et al., 2011; Lee et al., 2011). AtPRA1.B6 is localized to the ER and the Golgi (Jung et al., 2011), PRA1.F4 is found in the Golgi (Lee et al., 2017) whereas the subcellular localization of AtPRA1.B1 has not been demonstrated. SLY1, by acting in the ER and Golgi, could contribute to membrane fusion by interacting with Qa-SNAREs or nascent trans-SNARE complexes (Karnahl et al., 2018). The COPI coat composed of seven subunits (α/β/β’/γ/δ/ε/ζ) interacts with Golgi membranes (Aniento et al. 2022). The coatomer complex is not only involved in the biogenesis of COPI vesicles but it is also required to select the cargo to be included in the vesicles. Coat protein I (COPI) is necessary for intra-Golgi transport and retrograde transport from the Golgi back to the ER (Sánchez-Simarro et al., 2022). TRAPPC5 belongs to TRAPPI which functions in ER to Golgi transport (Vukašinović & Žárský, 2016). YKT61 is a unique R-SNARE lacking transmembrane domains (Bassham and Blatt 2008). Thus, it is present mainly in the cytoplasm and is critical for the dynamic biogenesis of vacuoles, for the maintenance of Golgi morphology, and for endocytosis, suggesting a broad role of YKT61-mediated vesicular trafficking in plant development (Ma et al., 2023). VSR1 is responsible for the sorting of proteins from the trans-Golgi network (TGN) to prevacuolar compartments (PVCs) and finally to their respective vacuoles (Shimada et al., 2003). The ESCRT machinery is responsible for the recruitment of the ubiquitinated cargo and membrane budding for ILV formation. Ubiquitinated cargoes are captured by ESCRT-0-like proteins, TOLs. The cargoes are subsequently translocated to the ESCRT-I, ESCRT-II and ESCRT-III (SNF7, VPS, ALIX) multiprotein complexes that constrict membranes to form intraluminal vesicles (Gao et al., 2017). The Qa-SNARE syntaxin SYP122 resides at the plasma membrane and mediates in the final stages of secretion (Waghmare et al., 2018).

**Table S1: Mass spectrometry proteomics data.**

**Table S2: XIC-based quantification of proteins identified in root insoluble proteomes.**

**Table S3: Proteins differentially accumulated in response to uranium (ANOVA test).**

**Table S4: Clustering of the 458 differentially accumulated proteins using the Self Organizing Tree Algorithm (SOTA).**

**Table S5: Main features of proteins differentially accumulated under U stress.**

**Table S6: Protein classification using PantherDB, QuickGO, and manual curation.**

