## Supplementary material for "Dynamics of the membrane- and cell wall-associated proteome of *Arabidopsis thaliana* roots in response to uranium stress": Supp Figures

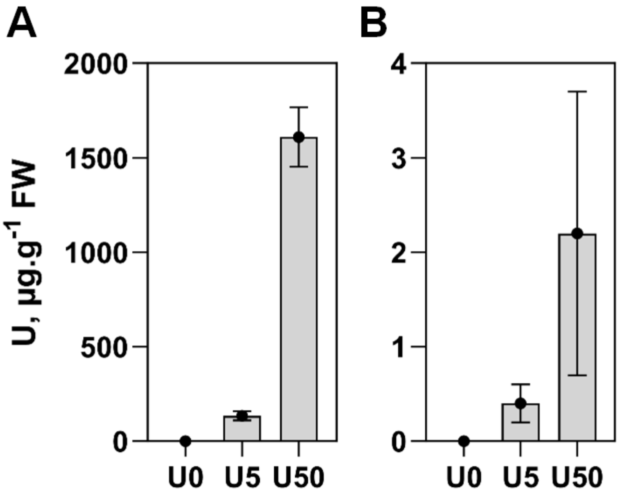


**Figure S1. Uranium content in roots and shoots of Arabidopsis plants.**
Uranium was measured by ICP-MS in roots (A) and shoots (B) of control (U0) and U-treated plants (U5 and U50 for 5 and 50 μM uranyl nitrate, respectively). Data are mean ± SD of n=6 biological replicates.

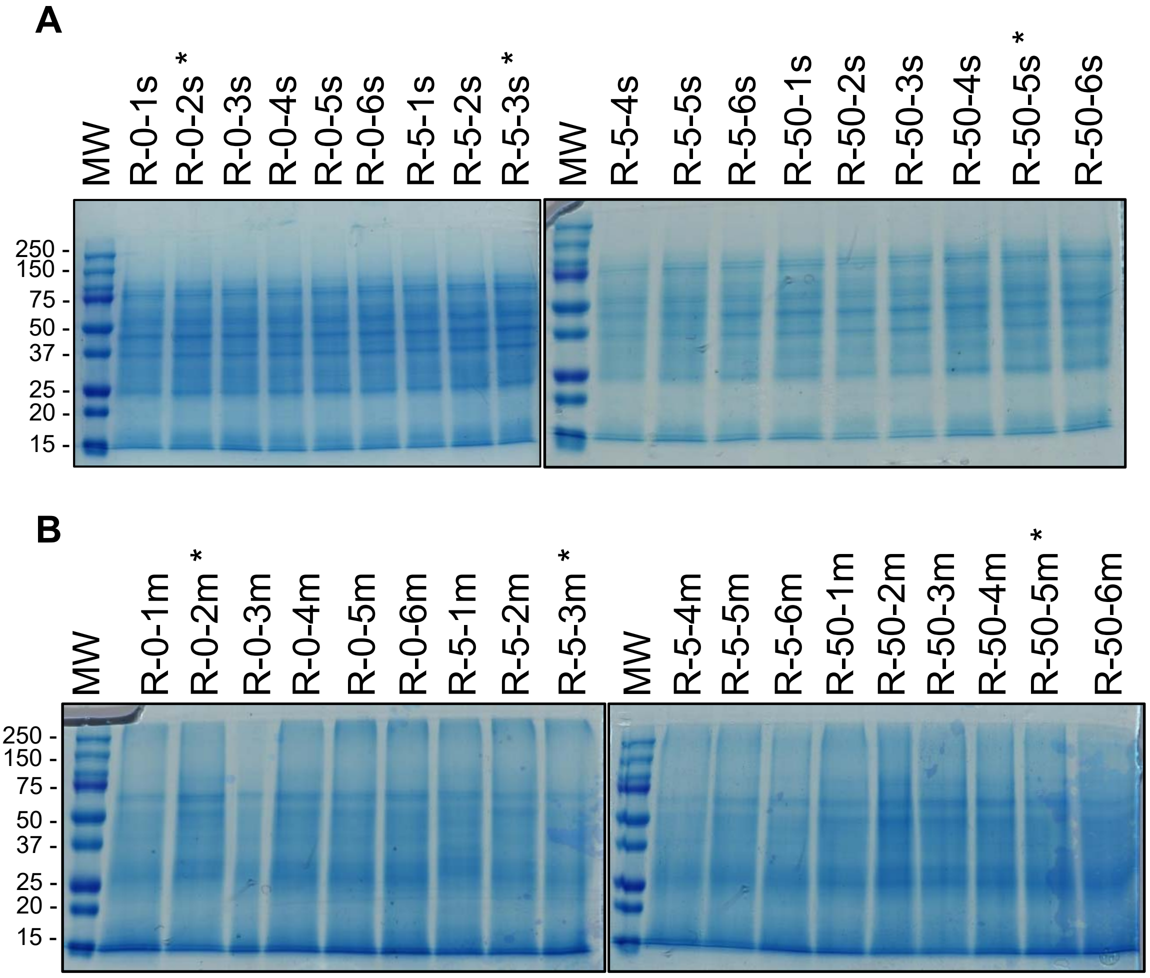


**Figure S2. SDS-PAGE analysis of soluble and membrane proteins from Arabidopsis roots.**
SDS-PAGE analysis of (**A**) soluble and (**B**) membrane proteins from control and U-treated *A. thaliana* roots. Proteins were stained with Coomassie Blue. Sample nomenclature: R, root; U concentration (0, 5 ,50 µM uranyle nitrate): 1s to 6s, biological replicates of soluble protein extracts; m1 to m6, biological replicates of membrane protein extracts; *, samples analyzed by western blot (Figure S3).

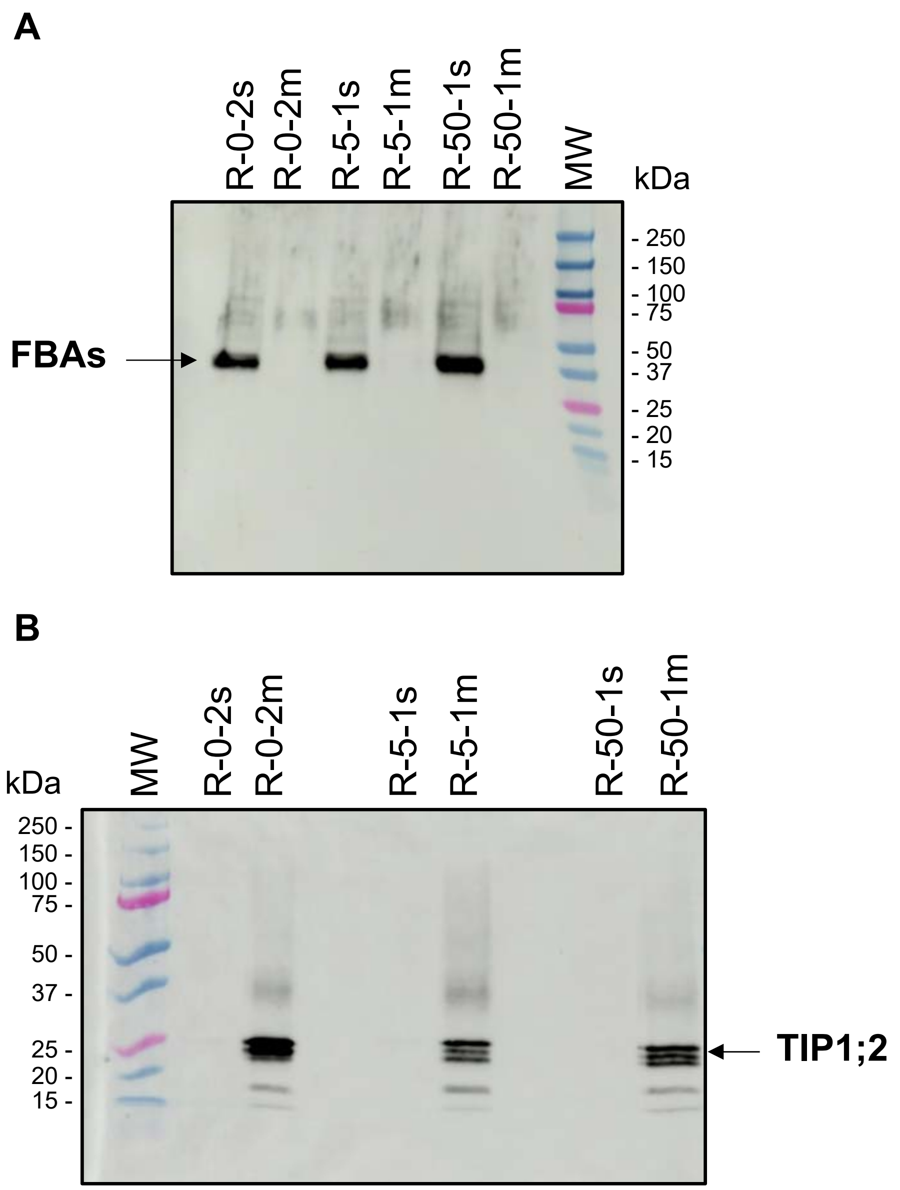


**Figure S3. Quality assessment of membrane proteins from Arabidopsis roots by Western blot analysis.**

Western blot detection of (**A**) the fructose-bisphosphate aldolases (FBAs) and (**B**) the tonoplastic intrinsic protein 1;2 (TIP1;2) in soluble (s) and membrane (m) protein extracts isolated from roots of Arabidopsis plants treated with 0, 5 and 50 µM uranyl nitrate. SDS-PAGE analysis of protein extracts (including sample nomenclature) is shown in Figure S2.

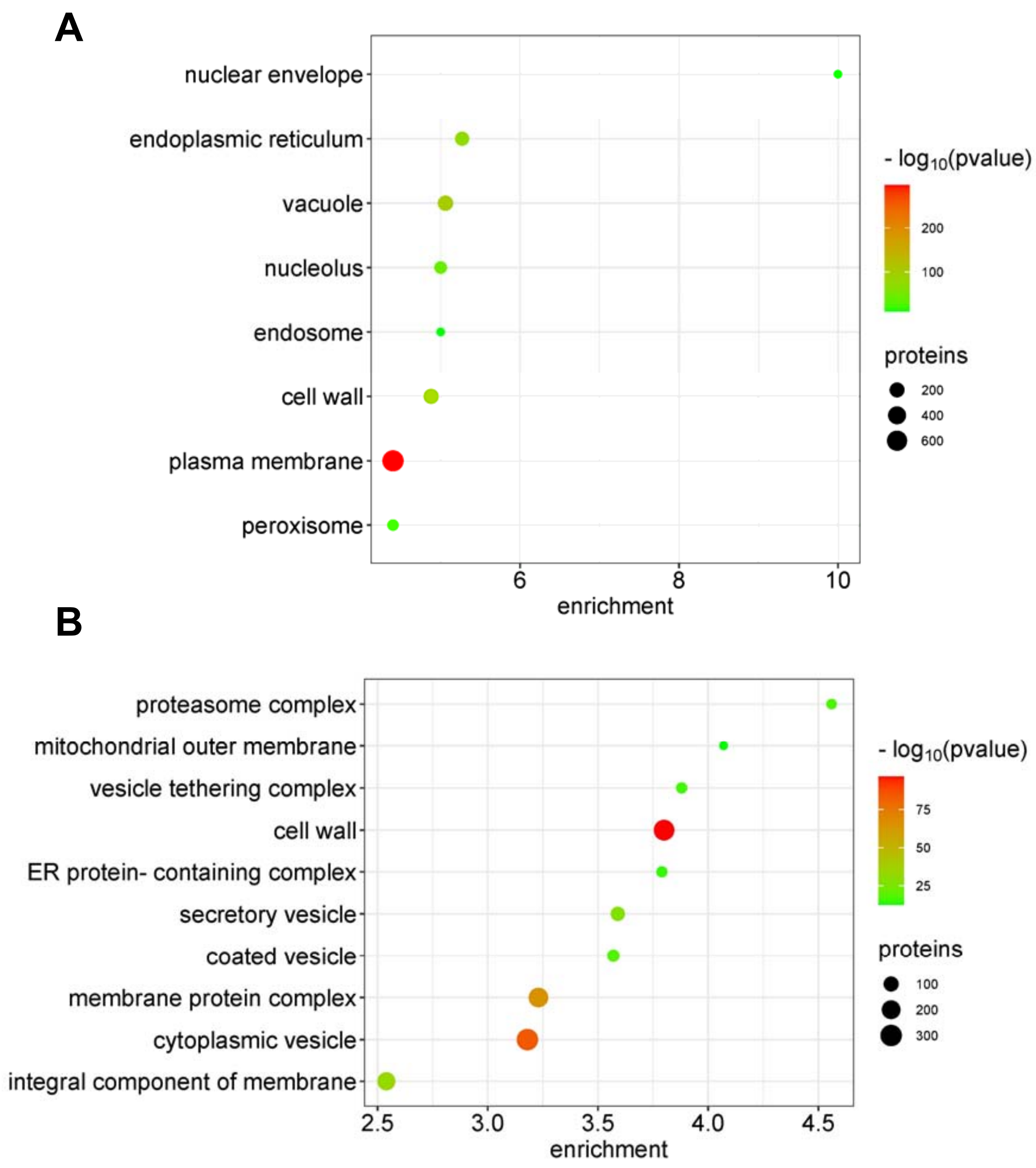


**Figure S4. GO enrichment analysis of cellular components in the membrane and cell wall proteome of Arabidopsis roots.**

GO enrichment analysis of cellular components was performed using the 2,802 proteins identified by mass spectrometry using the BiNGO (A) and Metascape (B) tools. Bubble plots show GO terms ordered by enrichment values (threshold >4 for BiNGO, >2.5 for Metascape). BiNGO settings to assess overrepresented GO cellular components were as follows: statistical hypergeometric test, Bonferroni Family-Wise Error rate multiple testing correction, and significant p-value <0.05. The Metascape enrichment analysis has been done with the GO cellular components ontology source. Terms with a p-value <0.01, a minimum count of 3, and an enrichment factor >2.0 have been grouped into clusters based on their membership similarities.

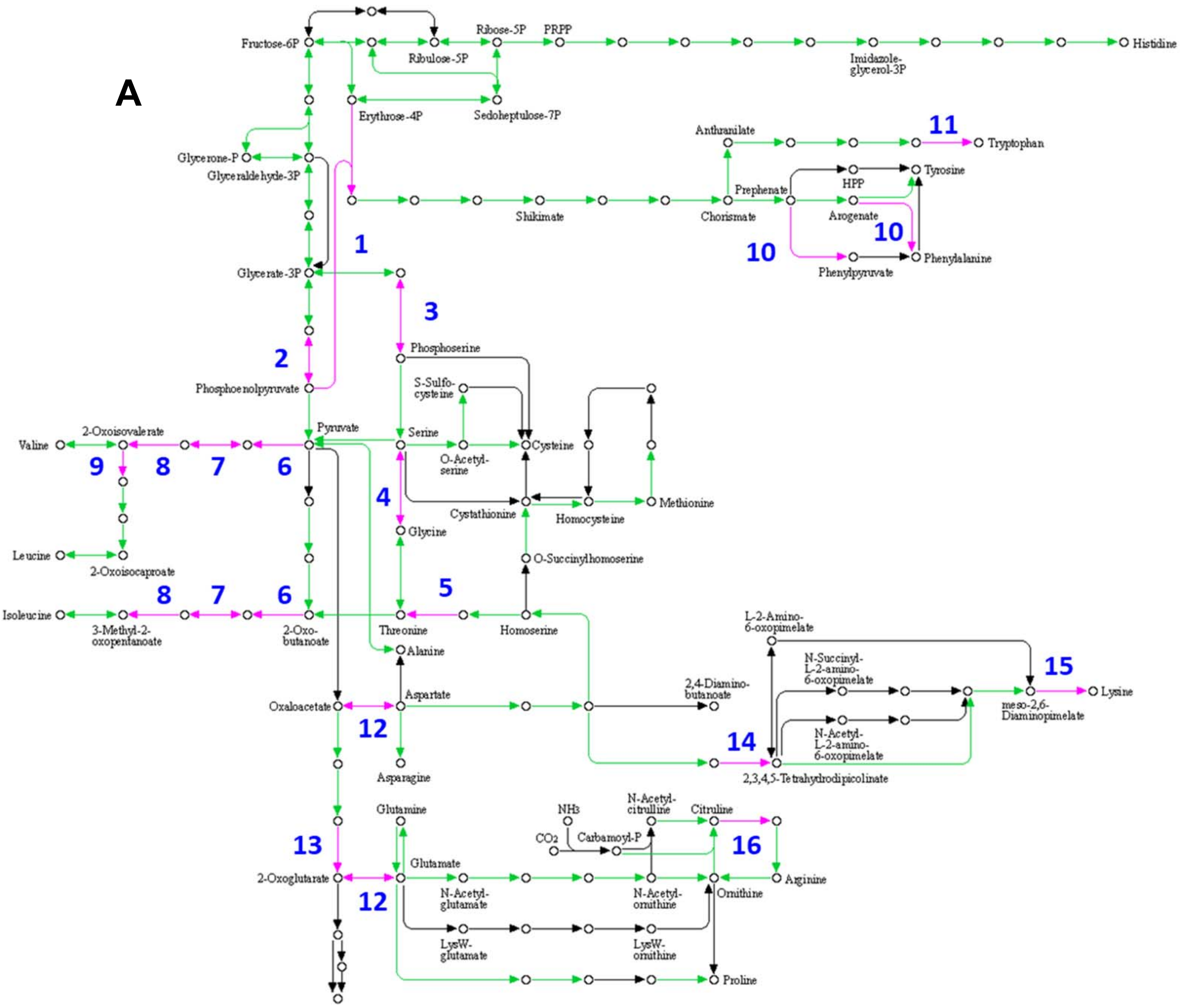


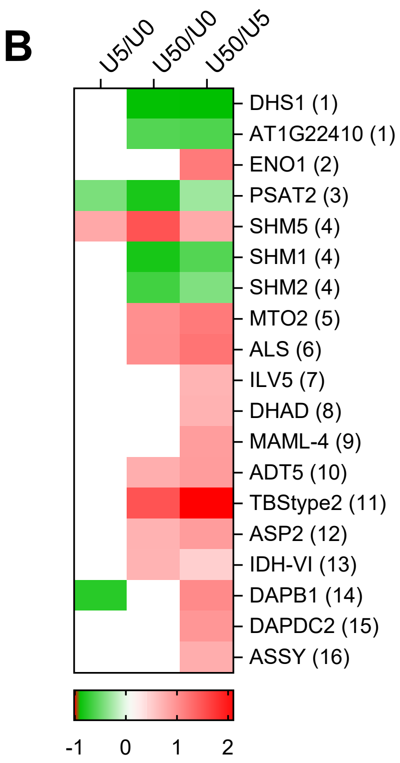


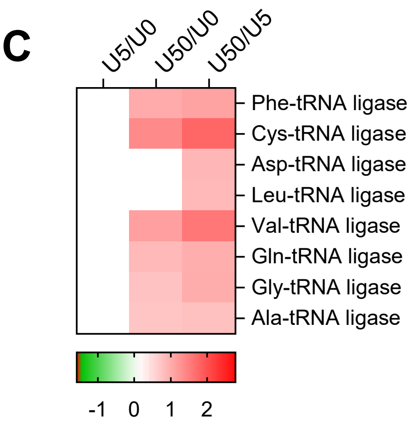


**Figure S5. Effect of uranium on amino acid metabolism.** (A) Proteins regulated by U are mapped to the KEGG pathway 'biosynthesis of amino acids' (ath01230). (B) Protein expression profiles are shown on a heatmap. The numbers in brackets refer to the enzyme positions in the pathway. (C) Heatmap of tRNA ligases regulated during U stress (not indicated on the pathway). Proteins whose abundance is significantly ($p<0.05$) increased or decreased in response to U stress are shown in red and green, respectively (log2 scale).

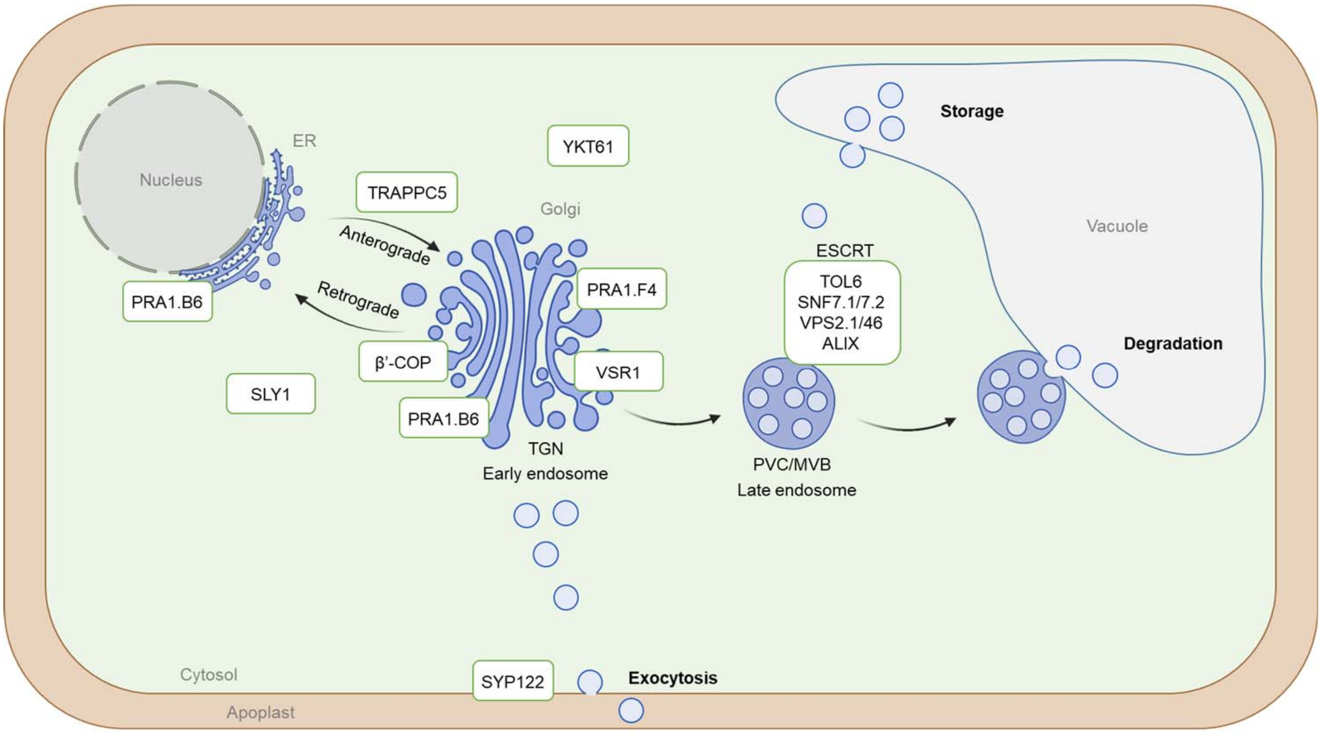


**Figure S6. Proteins differentially regulated by uranium involved in endomembrane trafficking.**

PRA1s play a role in the trafficking of cargo proteins destined to various endomembrane compartments (Jung et al., 2011; Lee et al., 2011). AtPRA1.B6 is localized to the ER and the Golgi (Jung et al., 2011), PRA1.F4 is found in the Golgi (Lee et al., 2017) whereas the subcellular localization of AtPRA1.B1 has not been demonstrated. SLY1, by acting in the ER and Golgi, could contribute to membrane fusion by interacting with Qa-SNAREs or nascent trans-SNARE complexes (Karnahl et al., 2018). The COPI coat composed of seven subunits (α/β/β'/γ/δ/ε/ζ) interacts with Golgi membranes (Aniento et al. 2022). The coatomer complex is not only involved in the biogenesis of COPI vesicles but it is also required to select the cargo to be included in the vesicles. Coat protein I (COPI) is necessary for intra-Golgi transport and retrograde transport from the Golgi back to the ER (Sánchez-Simarro et al., 2022). TRAPPC5 belongs to TRAPPI which functions in ER to Golgi transport (Vukašinović & Žárský, 2016). YKT61 is a unique R-SNARE lacking transmembrane domains (Bassham and Blatt 2008). Thus, it is present mainly in the cytoplasm and is critical for the dynamic biogenesis of vacuoles, for the maintenance of Golgi morphology, and for endocytosis, suggesting a broad role of YKT61-mediated vesicular trafficking in plant development (Ma et al., 2023). VSR1 is responsible for the sorting of proteins from the trans-Golgi network (TGN) to prevacuolar compartments (PVCs) and finally to their respective vacuoles (Shimada et al., 2003). The ESCRT machinery is responsible for the recruitment of the ubiquitinated cargo and membrane budding for ILV formation. Ubiquitinated cargoes are captured by ESCRT-0-like proteins, TOLs. The cargoes are subsequently translocated to the ESCRT-I, ESCRT-II and ESCRT-III (SNF7, VPS, ALIX) multiprotein complexes that constrict membranes to form intraluminal vesicles (Gao et al., 2017). The Qa-SNARE syntaxin SYP122 resides at the plasma membrane and mediates in the final stages of secretion (Waghmare et al., 2018).
